# Novelty and surprise-timing are broadcast by the basal forebrain

**DOI:** 10.1101/397513

**Authors:** Kaining Zhang, Charles D. Chen, Ilya E. Monosov

## Abstract

The basal forebrain (BF) is a principal source of modulation of the neocortex, and is thought to regulate cognitive functions such as attention, motivation, and learning by broadcasting information about the behavioral salience of events. An event can be salient because it is novel, surprising, or associated with reward prediction errors. But to date, the type of salience-related information the BF broadcasts is unclear. Here, we report that many BF neurons display phasic excitatory bursting that rapidly conveys the magnitude, probability, and timing of primary reinforcements. The same BF neurons also discriminate fully expected novel visual objects from familiar objects and respond to object-sequence violations, regardless of their relevance for subsequent behaviors, suggesting that they are not dedicated to signaling information about primary reinforcements. A different group of BF neurons displayed ramping activations that predicted the time of novel and surprising events. Their ramping was highly sensitive to the subjects’ confidence in event timing. Hence, BF neurons signal statistics about time and salience. Their activity may organize cortical computations to facilitate accurate behavioral responses to a diverse set of expected and ongoing events.

## Introduction

Theories of cognitive control predict that surprising and novel events, or events that deviate from our beliefs about the structure of the world, recruit attention and promote the formation of new memories. However, a clear biological understanding of the circuits that mediate surprise- and novelty-related behavior is missing. Particularly, how information such as timing of surprise and novelty are predicted and broadcast to influence sensory processing, learning, and cognition is not well-understood. One important challenge has been that surprise, novelty, value, and reward prediction errors are often difficult to distinguish and confounded (Wallis and Rich, 2011; Barto et al., 2013). For example, a surprising event is a deviation from the average of a range of possibilities (Hayden et al., 2011; Preuschoff et al., 2011; Barto et al., 2013) and can be experienced many times over. In contrast, a novel event is one that has not been experienced in the past (Anderson et al., 2008; Wang and Mitchell, 2011; Barto et al., 2013). Therefore, novelty arises from a comparison of current or ongoing events with representations of previous experiences. Theoretically novel events are always surprising, but surprising events are not always novel.

The basal forebrain (BF) is strongly implicated in modulation of attention, learning, and memory through gain modulation of ongoing event-related processing and plasticity (Morris et al., 1992; Voytko, 1996; Everitt and Robbins, 1997; Baxter and Chiba, 1999; Chudasama et al., 2004; Pinto et al., 2013; Avila and Lin, 2014; Peck and Salzman, 2014; Hangya et al., 2015; Lin et al., 2015; Raver and Lin, 2015). Previous work suggests that these BF functions may in part be mediated by single BF neurons that process anticipation and delivery of salient or surprising events. Two prominent neuronal activation patterns in the BF are observed following salient events: phasic bursting that has been identified in the brains of rodents (Lin and Nicolelis, 2008; Hangya et al., 2015) and tonic activations (Hangya et al., 2015; Monosov et al., 2015), which in monkeys are often seen in neurons that also ramp to the time of delivery of uncertain or aversive outcomes (Monosov et al., 2015).

To date, it remains unclear how these neuronal activations signal surprise and/or novelty, and how their surprise-related responses relate to errors in the animals’ estimates of state values, often referred to as reward prediction errors (or RPEs). Therefore, how BF activations contribute to cognitive function remains unclear.

To answer these questions, we sought to 1) confirm the existence of phasic burst neurons in monkeys (so far, they have explicitly only been demonstrated in rodents), 2) assess whether prediction-related phasic bursting and ramping activity occurs in distinct groups of neurons, and 3) study BF neurons while monkeys participate in a series of behavioral procedures that test whether and how BF represents prediction errors, surprise, value, novelty, and timing.

## Results

### Phasic and ramping activity is observed in distinct BF cell groups that differentially encode reinforcement statistics

The first experiment aimed to 1) identify phasic bursting neurons in monkeys, 2) assess if they are a distinct functional class of neurons from the previously identified BF ramping neurons, and 3) study how phasic bursting and ramping neurons signal predictions about reinforcement.

We recorded the activity of BF neurons (Methods) while monkeys participated in a Pavlovian procedure in which they experienced predictions of rewards that varied in magnitude and probability (Monosov and Hikosaka, 2013; Monosov et al., 2015; White and Monosov, 2016; Monosov, 2017). A reward-probability block contained five visual conditioned stimuli (CSs) associated with five probabilistic reward predictions (0, 25, 50, 75 and 100% of 0.25ml of juice). A reward-amount block contained five other CSs associated with certain reward predictions of varying reward amounts (0.25, 0.1875, 0.125, 0.065 and 0ml). The expected values (EVs) of the five CSs in the probability block matched the expected values of the five CSs in the amount block. The two-block design removed confounds introduced by risk seeking-related changes in subjective values of the CSs (Monosov and Hikosaka, 2013; White and Monosov, 2016; Monosov, 2017). After conditioning (Methods), we recorded the activity of BF neurons in 6 monkeys. Any neuron that displayed ramping and/or phasic burst responses to any of the task events in the Pavlovian behavioral procedure was recorded (n=70).

Two example BF neurons are shown in Figure 1A. The first neuron (Figure 1A – top) displayed short latency bursting after the presentation of the probability and amount CSs. This phasic activation was greatest following the presentation of the CS associated with the highest EV, in either the reward probability or the reward amount block, and least following the presentation of the CSs associated with no reward. Hence, in either block, the bursting activity was strongly correlated with expected value (Probability block, Spearman’s rank correlation ρ = 0.84, p < 0.0001, Amount block, ρ = 0.86, p < 0.0001). The second neuron (Figure 1A – bottom) had a very different response. Shortly after the CSs were presented, it displayed a consistent CS-onset related inhibition that was greatest in the low value trials and less apparent during high value trials. In the reward probability block, this initial change was followed by ramping activity to the time of the risky reward delivery (following 75, 50, and 25% CSs). The neuron’s activity was better fit by a model of uncertainty (ρ = 0.77, p = 0.0001, measured in last 500ms) than EV (ρ =-0.01, p = 0.94). In the reward amount block, in which all trials were certain, the neuron signaled EV until the time of the reinforcement by its tonic activity (ρ = 0.70, p < 0.0001, last 500ms). These two example neurons highlight the possibility that the monkey BF may contain distinct classes of neurons: phasic bursting neurons that co-vary with the magnitude and probability of reinforcements and tonic neurons that display ramping activity that predict the timing of uncertain outcomes.

**Figure 1:**
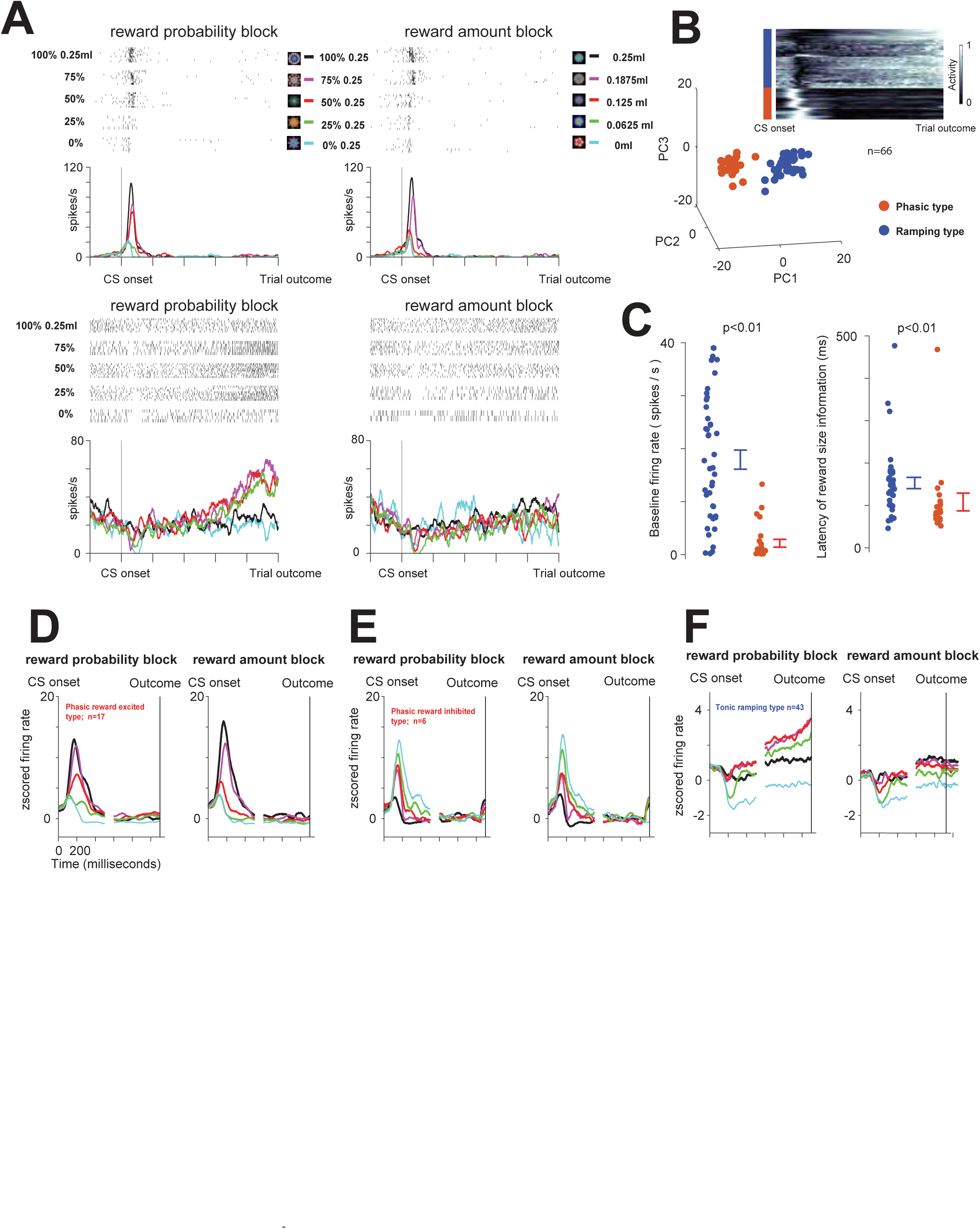
Two groups of BF neurons encode the magnitude and probability of reinforcement in distinct manners. (**A**) Responses of two example BF neurons (top and bottom) to the presentation of 10 fractal objects associated with certain and uncertain predictions of juice reward in the reward probability block (left) and reward amount block (right).(**B**) Clustering of BF neurons based on average activity in the probability block. Inset heat map shows the activity of 66 BF neurons (normalized from 0 to 1) from the time of the CS onset to the time of the trial outcome (reward or no reward). Each line represents the average activity across all 5 trial types in the probability block for each neuron. Below are the results of PCA analyses performed on those normalized CS response functions. K-means clustering (Methods) was used to separate the neurons into two groups: red group (n=23) and blue group (n=43). The group identities of the neurons are also indicated by a color bar to the left of the heat map. (**C**) The two clusters of neurons (red and blue) display distinct baseline firing rates (left) and latencies of value coding in the reward amount block (right). Each dot represents data from a single neuron. Error bars around the mean show SEM. (**D**) Average responses of the neurons in the blue group in the reward-probability block (left) and reward amount block (right). (**E-F**) Average responses of the neurons in the red group in the reward-probability block (left) and reward amount block (right). **E** shows neurons that displayed greater activation for reward versus no reward trials, while **F** shows neurons that displayed greater activation for no reward trials.

To test whether this was the case, we clustered BF neurons based on their average response vectors. Only the neurons that had been recorded in every condition in both blocks were included (n=66/70). Importantly, neuronal response vectors were obtained by averaging the neuronal activity across all five CSs in the reward-probability block, and were subsequently normalized (from 0 to 1; Figure 1B; inset). In this way, neuronal tuning (e.g. representation of reward probability) and baseline firing rates were not considered in the clustering analysis.

Principal component analyses clustering revealed two distinct clusters (Figure 1B; Methods). The cluster identity of each single neuron’s response vector is color coded in Figure 1B-inset. The first cluster (*red; n=23*) showed clear bursting after the CS onset (see single neurons response vectors in Figure 1B inset). In contrast, the second cluster (*blue; n=43*) showed an initial suppression following the CS onset, and slow ramp-like increase in activity as the trial’s outcome neared.

These two clusters had different baseline firing rates (Figure 1C): one had relatively high firing rates (blue cluster; average frequency = 18 Hz; S.D. = 12 Hz) and the other had low firing rates (red cluster; average frequency = 2.1 Hz; S.D.= 3.5 Hz). In addition, we found that both clusters’ initial CS responses co-varied with the magnitude of predicted reward, but the timing or latency of this information was different among them. Reward size information was conveyed earlier by the neurons in the phasic bursting red cluster (Figure 1C-right; rank sum test; p<0.01; blue cluster, average = 157 ms, S.D. = 78 ms; red cluster, average = 125 ms, S.D. = 103 ms). Also, the two clusters of neurons differed in how they represented probability and amount of reinforcement (Figure 1D-F; Supplemental Figure 1). Phasic bursting neurons (*red cluster*) signaled the expected values of the CSs in their bursting activations. The bursting activity was correlated with the probability in the probability block (ρ = 0.60, p<0.0001) and reward amount in the amount block (ρ = 0.71, p<0.0001). The tonic ramping neurons’ initial suppression co-varied with expected value (ρ = 0.47, p < 0.0001 in probability block, ρ = 0.46, p <0.0001 in amount block). But in trials in which reward was uncertain (or risky) they also displayed ramping activity to the trial outcome as previously reported (Monosov et al., 2015). This activity was correlated with uncertainty (ρ = 0.71, p < 0.0001). The locations of the phasic bursting and tonic ramping neurons were reconstructed in Supplemental Figure 2 and confirmed using in-vivo MRI methods (Supplemental Figure 3). Both phasic bursting and tonic ramping neurons were found within the BF areas that overlap with the diagonal band of Broca and Nucleus Basalis of Meynert (Mesulam et al., 1983; Monosov et al., 2015; Turchi et al., 2018).

**Figure 2:**
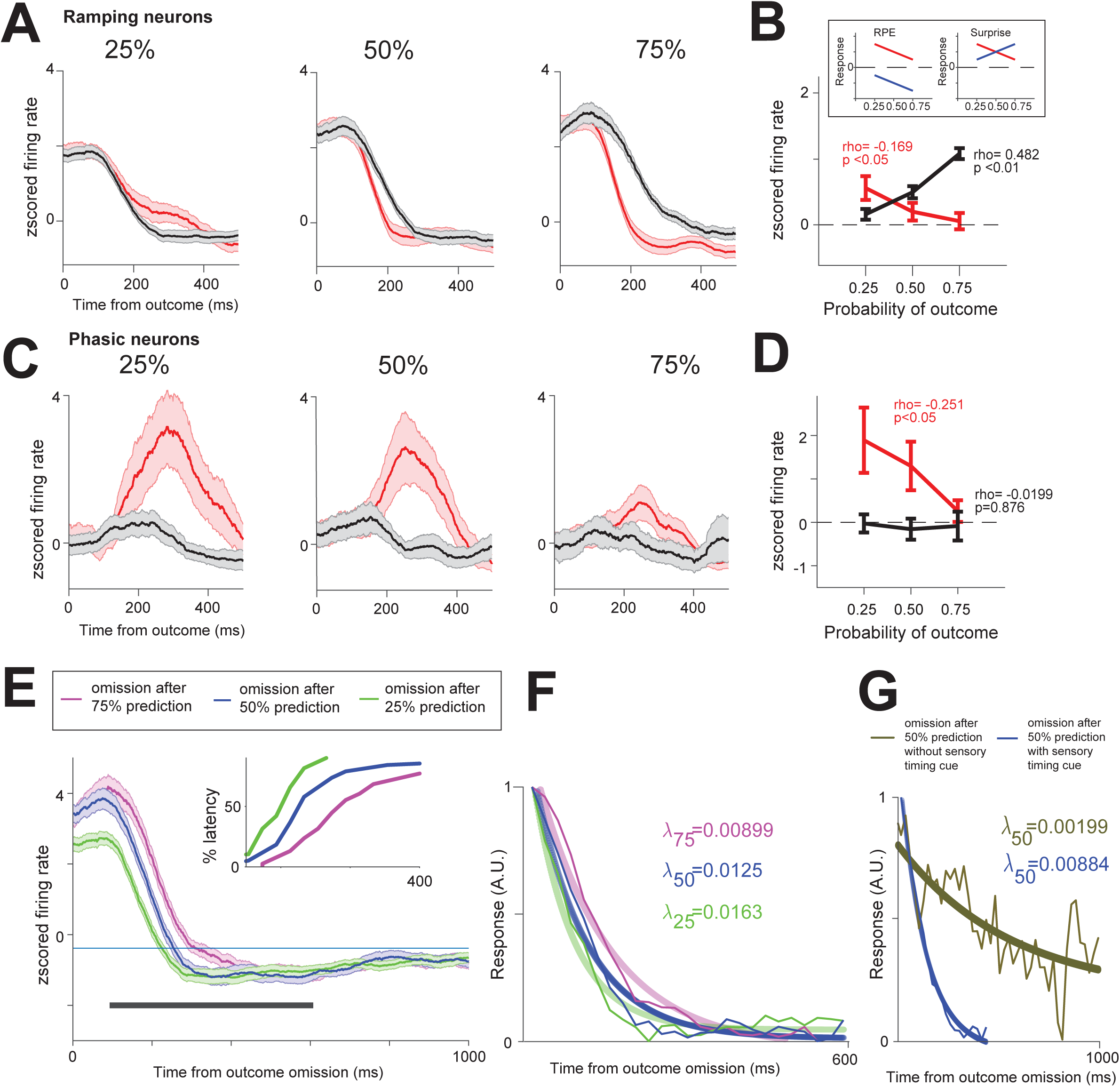
Differential coding of surprise in ramping and bursting BF neurons. (**A**) Ramping neurons’ average outcome activity in 25%, 50% and 75% conditions. Red – reward delivered trials; black – no reward trials. (**B**) Ramping neurons’ average responses for reward delivery and no reward trials. Linear correlations of responses with reward expectancy are indicated (time window: 100ms to 400ms). Inset shows cartoon models of reward prediction error coding (left) and reward surprise coding (right). BF ramping neurons’ responses resembled surprise. (**C**) Outcome activity of phasic bursting neurons. Conventions are the same as in (A). (**D**) Phasic bursting neurons’ responses during resembled reward prediction error coding only in reward delivery trials (red; time window: 200ms to 500ms). (**E**) Activity of BF ramping neurons during 25, 50, and 75% reward probability trials in which the reward was omitted. The ramping activity returned to inter-trial baseline level (thin blue line) at different latencies across these three types of trials: earliest during 25% trials, and latest during 75% trials. Cumulative distributions of these latencies are shown in the inset. (**F**) Exponential fits (thick lines) to the population binned activity (thin lines; Methods). Fits and decay rates (right) were calculated for the population after the activity for each trial type was normalized from 0 to 1, such that for each of the tree conditions, the starting point is 1. (**G**) Same as **F,** except here we compared fit and decay rate during 50% trials in which an explicit cue indicated the end of the trial (dark blue) with fit and decay rate during 50% trials in which no explicit cue was given (and the CS remained on the screen; Methods).

**Figure 3:**
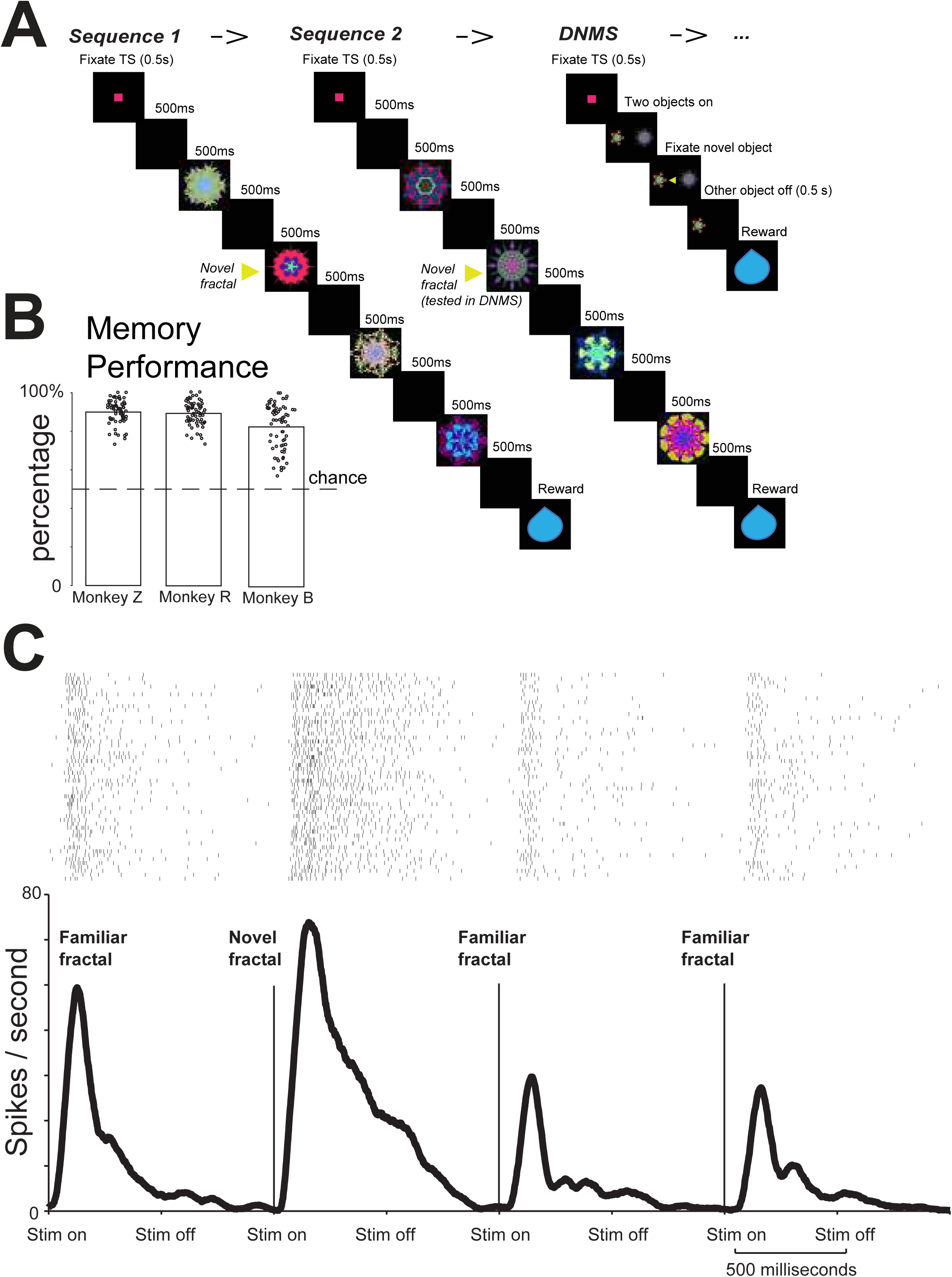
Object sequence task. (**A**) The monkey was first shown two sequences of fractals. Each sequence contained 4 fractals, in which 1^st^, 3^rd^, 4^th^ fractals were fixed familiar objects and 2^nd^ fractal was always novel. After the two sequences, the monkeys performed a delayed non-match to sample task (DNMS) in which one object was novel and the other was the object that was previously novel in sequence 2. Monkeys fixated the novel object for reward. (**B**) Behavioral performance for three monkeys. Y-axis shows the percentage of first saccades to the novel object in DNMS, the percentages are significantly different from 0.5 for all three monkeys (p<0.01, sign-rank test). (**C**) An example BF phasic bursting neuron’s responses to the four objects in in a sequence. The response was highest for the second (novel) fractal (rank sum test; p<0.05).

These data show that the primate BF contains roughly distinct groups of neurons that exhibit bursting and ramping responses following reinforcement predictions, and that the short latency bursting in the BF conveys higher-order information, such as information about both reward probability and amount at a very short latency (Figure 1C-right).

### Phasic and ramping neurons differentially encode early versus late rewards under temporal uncertainty

Several important questions arose from the data in Figure 1. First, might the ramping of BF neurons encode the estimated time of uncertain rewards? If so, then if rewards were certain but their timing was uncertain, they should display ramping activity to the time of the earliest possible reward. Second, phasic bursting neurons bursts seemed to scale with the value of the CS, regardless of whether the value was manipulated by changing probability or amount of reinforcement. Might these neurons also encode the subjective value of early versus late rewards?

To answer these questions, we designed a reward timing procedure (Methods). Here, five distinct visual fractal objects serve as conditioned stimuli (CSs) that predict either (1) a probabilistic delay before a reward with deterministic delivery (reward-timing-uncertain CSs); or (2) a deterministic delay before a reward with some probability of delivery (reward-probability CS; Figure 1A). In trials with one of the four reward-timing-uncertain CSs, reward was always delivered either 1.5 s after CS onset or 4.5 s after CS onset. Depending on the reward-timing-uncertain CS, the probability that a reward was delivered 1.5 s after CS onset was 0.25, 0.50, 0.75, or 1 (25, 50, 75, 100%; Supplemental Figure 4). In trials with the reward-probability CS, reward was delivered with a delay of 1.5 s after CS onset with 0.50 probability. Monkeys’ licking behavior suggested that they were sensitive to the timing and probability of reward (Supplemental Figure 4B). First, during the four reward-timing-uncertain CSs, monkeys displayed increased licking behavior before 1.5 s, then a decrease in licking behavior after 1.5 s if reward was not delivered, then finally an increase in licking behavior to the time of reward at 4.5 s. During reward omissions, in 75% reward trials licking behavior remained higher than 25% and 50% trials, even 0.5 s after the short delivery time (p <0.01, rank-sum test, time window 2s to 2.5s after the onset of fractal), indicating that the monkeys’ confidence in the outcome influenced their anticipatory behavior. Finally, the mean magnitude of anticipatory licking behavior before possible reward delivery at 1.5 s across all trials increased with the probability of reward delivery at 1.5 s (Spearman’s rank correlation, ρ=0.38, p=<0.0001; Supplemental Figure 4C).

**Figure 4:**
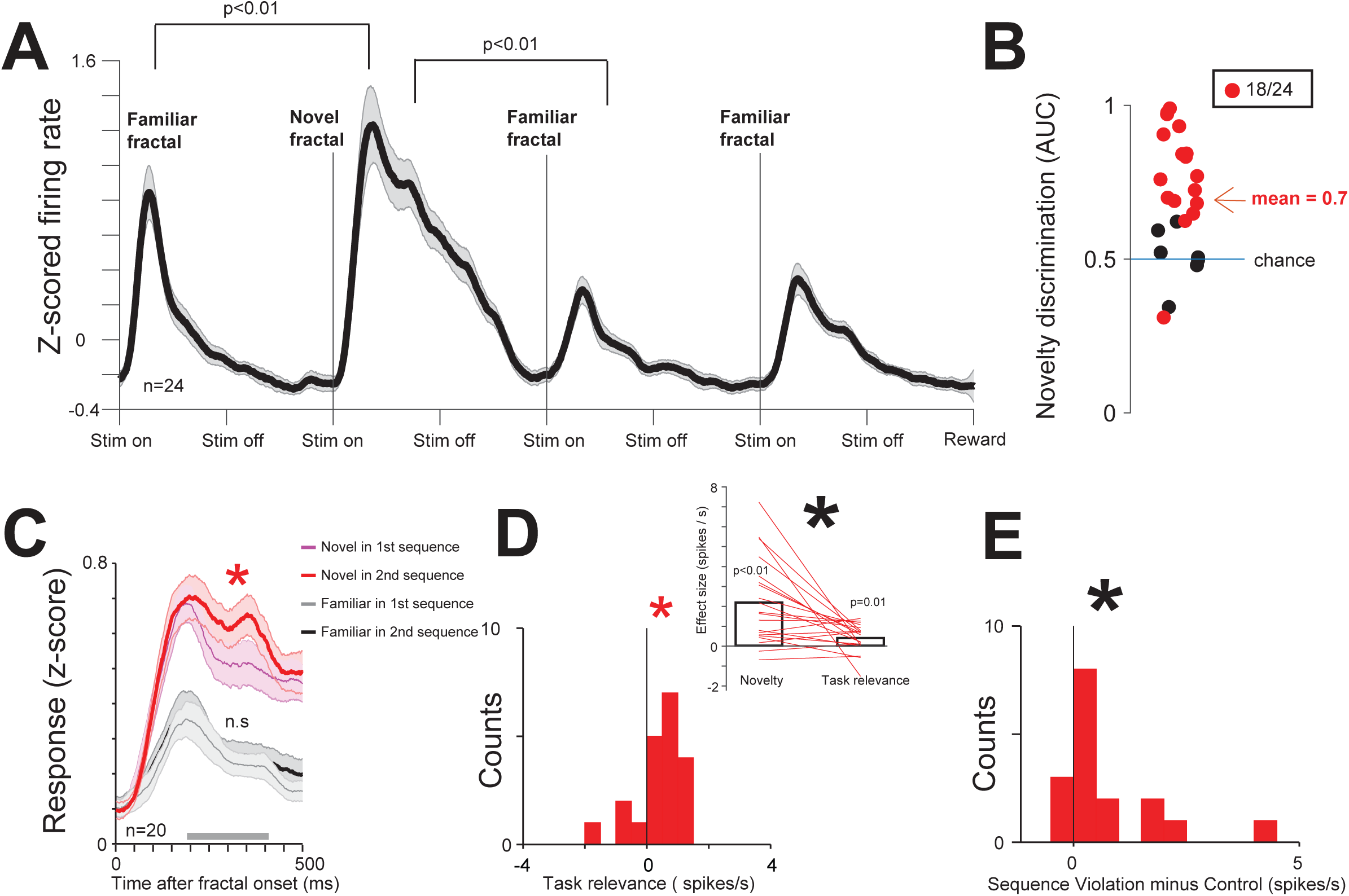
Phasic bursting neurons signal novelty and surprise not directly related to reward. (**A**) Average activity of phasic bursting neurons in the object sequence task. (**B**) Area under the ROC curve (AUC) for each phasic neuron that assessed the ability of the neuron to discriminate novel versus familiar objects. Red dots are neurons that can significantly discriminate novel versus familiar objects (time window: 200 ms to 400 ms). (**C**) Phasic neuron’s group average responses to novel fractals in Sequence 1 (thin blue line), Sequence 2 (thick red line), and the to the last 2 familiar fractals in Sequence 1 (thin gray line) and Sequence 2 (thick black line). Shaded region represent SEM. Asterisk indicates significant difference (**p<0.05**) between novel fractal responses in Sequence 1 and 2. (**D**) Lower left: Histogram of single neurons’ response differences for novel fractal in Sequence 2 and Sequence 1. Red asterisk indicates significant difference from 0 (p<0.05), Black asterisk indicates significant difference from 0 (p<0.01). Upper right: for each neuron, the data from the histogram (on the right) compared with the strength of novelty discrimination (left; defined as the difference between neuronal responses for X and Y). Both novelty discrimination and task-relevance effects are significant, but the novelty effect is stronger (p<0.05). (**E**) At low probability (11 %) one of the familiar fractal in sequence 2 (or 4) was substituted by a familiar fractal (Methods). Phasic neurons’ responses were enhanced (p<0.01) by this object sequence violation.

To test how phasic bursting neurons and tonic ramping neurons encode temporally-uncertain reward predictions, we first identified them using the task in Figure 1, and then recorded them in the reward timing procedure.

Tonic ramping neurons displayed ramping activity in the reward-timing uncertain conditions (0.75, 0.5, 0.25; Supplemental Figure 5A) and this activity was correlated with the probability of reinforcement at 1.5 s (Spearman’s rank correlation, ρ=0.61, p<0.0001, analysis window, 1s to 1.5s). Similar results were obtained in the probability-amount task (Figure 1). In trials in which ramping activity was observed (75, 50, 25%), the activation of this class of neurons was correlated with the probability of reinforcement (Spearman’s rank correlation, ρ=0.2, p=0.025, analysis window, 0.5s before outcome is resolved).

When reward was omitted at 1.5 s, the activity also displayed ramping to the late reward at 4.5 s (Supplemental Figure 6-top). Therefore, tonic ramping BF activity tracks reward delivery during temporal reward uncertainty (before 1.5 s) and during long epochs (in which there is uncertainty due to noise in interval timing).

On the other hand, phasic bursting neurons’ activity scaled with reward timing such that highest activity was evoked by CSs predicting the earliest reward (Supplemental Figures 5-6). Their average activity was correlated with reward probability at 1.5 s (Spearman’s rank correlation, ρ=0.55, p<0.0001, analysis window, 0.1s to 0.6s).

To summarize, in the monkey BF, a population of phasically active bursting neurons can rapidly broadcast higher-order information about salient reinforcements, including information about magnitude, probability or timing. Also, the monkey BF contains a population of tonic ramping neurons that signal similar information in their tonic activity (that lasts through the entire CS epoch). These neurons also encode the estimated timing of salient uncertain events (which could be uncertain due to errors in timing or due to the probabilistic nature of the outcome). Their ramping activity is correlated with the monkeys’ confidence in obtaining reward at the time of the trial outcome (i.e., 75>50>25%).

### Basal forebrain phasic and ramping neurons signal reinforcement surprise in distinct manners

A long-standing question is whether the BF signals errors in state values, *reward prediction errors* (RPEs) – a key signal for updating reward values and mediating economic choice (Schultz et al., 1997; Schultz, 2002; Lak et al., 2014). An alternative hypothesis is that BF neurons signal a rectified or an unsigned prediction error rather than a value (signed) prediction error. To differentiate unsigned RPEs from signed (‘reward value’) RPEs, it is important to assess neuronal responses to reward deliveries and reward omissions after 25, 50, and 75% predictions (e.g., in many instances 50% is not sufficient; see toy models in Supplemental Figure 7).

If neurons in the BF signal unsigned value of reward prediction errors, then BF neurons should display greatest responses to reward deliveries following 25% reward predictions, and smallest responses following a 75% reward prediction. The same neurons should display greatest responses to reward omissions following 75% reward predictions, and smallest responses following 25% reward predictions. An alternative possibility is that BF neurons encode reward omissions in a signed manner (consistent with the signed RPE hypothesis). In that case, BF neurons may display inhibitions following omissions that are inversely related to the probability of reward.

We found that BF ramping neurons outcome related activity was correlated with unsigned prediction errors (Figures 2A-B). After the trial outcome, the magnitude of their activity was greatest during reward-delivered trials following 25% reward predictions, and greatest during reward-omission trials following 75% reward predictions. Reward omission and reward delivery outcome responses were significantly correlated with expectancy (Figure 2B), albeit in opposite manners.

BF phasic bursting neurons’ outcome related activity signaled prediction errors following deliveries. Their delivery responses were correlated with expectancy (Figure 2D), displaying highest activations following reward deliveries in 25% reward trials. However, unlike the ramping neurons, these neurons did not discriminate reward omissions following different uncertain reward predictions (Figures 2C-D; black).

Surprise has a temporal dimension, and BF ramping neurons clearly display ramping signals (Figures 1-2), which are correlated with the monkeys’ confidence in reward delivery, and which may be ideal encoding estimates of outcome timing. If this is the case, the ramping-down activity following outcome delivery or omission should be partly mediated by the subjects’ confidence.

To test this, we took advantage of the fact that in our task the CSs co-terminated with the reinforcement deliveries (both rewards and omissions), and no other external cues indicated that an omission has occurred. If ramping activity in part reflects the animals’ temporal estimates, then we should see different ramping-down responses following omissions in 25, 50, and 75% trials. This is indeed what we found. Following reward omission, the BF ramping activity returned to baseline earliest during 25% reward trials and latest during 75% trials. The decay of the ramping also roughly scaled with reward expectation: decay rate was greatest following omissions during 25% and least during 75% trials (Figure 2F; the 95% confidence intervals of 25%, 50% and 75% decay rates exclude each other; Methods). Note that different firing rates across trial conditions could not explain these results because to calculate the decay, we first normalized each trial type from 0 to 1 (Figure 2F; Methods).

To further test the hypothesis that the BF ramping activity reflects timing estimates, we studied the activity of BF ramping neurons in the reward timing procedure because it contained two distinct 50% reward predictions: one in which the CS co-terminated with the outcome and one in which the CS remained on the screen (Supplemental Figure 4-5). In the first condition, the animals obtained a precise signal about the timing of the trial (e.g. when to expect a delivery or an omission) while in the other they did not. We again found that the decay rate of BF ramping neurons was sensitive to temporal predictions: it was significantly greater when animals did not receive an explicit temporal cue (Figure 2G; the 99% confidence intervals of the decay rates exclude each other; Methods). These data show that BF ramping activity is strongly influenced by evidence and confidence about the timing of reinforcements.

### Phasic and ramping BF neurons differentially represent object novelty and sensory surprise

The data thus far show that BF neurons are sensitive to surprise. Ramping neurons signal surprise about internal and external events and are sensitive to estimates of timing. Phasic bursting neurons signal surprise about deliveries (external events) and are not consistently tuned to value-differences of reward omissions. However, surprises arise due to violations in belief states following a probabilistic prediction, when there is a deviation of the outcome from the mean of expected outcomes (Barto et al., 2013), or as a result of novelty due to a comparison of ongoing sensory events with representations of past experiences. Next, we studied if and how different sensory surprises are encoded in BF.

To test how BF represents novelty we designed a behavioral procedure in which novelty was fully expected. Here, novelty was surprising, but not due to a deviation from the animals’ expectations in its occurrence.

Monkeys experienced four distinct sequences of object presentations (S1, S2, S3, S4). Each sequence contained 3 familiar objects and 1 novel object. The novel object was always presented at the same point in the sequence (Figure 3A). If a neuron has a selective novelty response, it should respond more strongly and consistently to the novel object than to the familiar objects in the sequence. To assess if novelty-responses were dominantly due to task-relevance or reward-prediction, following S2 and S4, monkeys performed a reaction-time Delayed-Non-Matching-to-Sample task (DNMS, Figure 3A-right). During DNMS, an object that was novel during the presentation of S2 (or S4 if the DNMS trial followed S4) was presented with a novel object that has never been experienced. The trial continued until the monkeys fixated the novel object for 0.5 second to get reward (Figure 3A – right). The monkeys were never penalized for looking at the previously experienced object. Thus, the novel objects in S2 or S4 did not have an explicit reward association, but aided the monkey in subsequent DNMS trials. Monkeys’ behaviors indicated that they understood the task and utilized previous experiences to increase their reward rate. Their first saccade following the presentation of the two fractals most often landed on the novel object and remained there until the other stimulus disappeared and reward was delivered.

A key finding was that phasic BF phasic bursting neurons robustly discriminated the novel object from the familiar objects. An example BF phasic bursting neuron is shown in Figure 3C. This neuron responded selectively to the novel object (p<0.01; rank sum test). This selective response could not be explained by priming or reward-proximity because the novel objects always appeared in the second position in the sequence (Figure 3A) rather than the first or the last. Like the example neuron, the population BF phasic bursting response (Figure 4A) and the single neurons (Figure 4B) selectively discriminated the novel object versus familiar objects.

Importantly, BF phasic neurons’ responses were activated by novel objects regardless of their task relevance (Figure 4C). That is, during both S1 and S3, BF phasic neurons displayed stronger responses to novel objects than to familiar objects (independent sign rank sum tests, p<0.01). However, their novelty responses were also modulated by task relevance. A small but highly consistent increase in firing was observed for novel objects shown in S2 and S4 (Figure 4C-D) versus novel objects shown in S1 and S3. This could not be explained by a general arousal increase in S2 and S4 relative to S1 and S3 because the activity for the third and fourth familiar fractals in the sequences was not different across S2 and S4 versus S1 and S3 (Figure C).

In contrast to the phasic BF neurons, the ramping neurons did not discriminate novel from familiar objects (Supplemental Figure 8). However, they did display ramping activity that anticipated two critical events in the object sequence task: novel object presentation and reward (Supplemental Figure 8). Because of this finding, and our previous work that indicated that these same neurons also display ramping activity to the estimated time of punishment delivery (Monosov et al., 2015), we asked whether they would also ramp to salient distracting sensory events (e.g. bursts of noise) and found that they did not (Supplemental Figure 9). These data are consistent with our previous observations that this population of BF neurons ramp to the estimated timing of salient events, such as novel objects, and reinforcements, particularly when they are expected following a long delay (Supplemental Figure 6).

Finally, we assessed how BF neurons signal surprises that arise due to errors in object order in the object sequence task. Previous work showed that the temporal cortex, a major target of BF projections (Mesulam et al., 1983), is sensitive to sequence-violations (Meyer and Olson, 2011). Might BF be a candidate source region for such a signal in the cortex? To study this, we replaced an object in the S2 with an object from S1, or an object from S4 with an object from S3 on approximately 11% of trials. Importantly, these replacements avoided reward prediction errors because during replacements, the proximity to reward was not changed (e.g. a fractal that normally appears in the third position in in S1 could appear in the third position in S2). Using this manipulation, we found that object sequence violations produce small but significant increase in the population responses of both phasic and tonic BF neurons (Figure 4 and Supplemental Figure 8). These data indicate that the BF can broadcast information about novel and surprising sensory events that are not directly associated with primary reinforcements, such as rewards and punishments.

## Discussion

Natural environments demand that humans and other animals detect and act on salient, surprising, and novel stimuli. However, how the neocortex detects and preferentially processes novelty and surprise in a temporally precise manner are poorly understood. BF is one of the principal sources of modulation and control of the neocortex. However, few studies have examined the events and behavioral states that elicit activity in the primate BF. Here we studied two types of BF activations: phasic bursting and tonic ramping. We found that these two types of activations are found in two separate groups of neurons and that these populations of neurons distinctly broadcast information about surprise, time, and novelty.

The first group of neurons displayed stimulus evoked phasic bursting that scaled with reward expectancy and value. Their CS-evoked bursting was greatest when the value of a CS was relatively large because the CS was associated with a large reward, a highly probable reward, or an early reward. Their trial outcome responses of these neurons were also strongly stimulus and expectancy driven. Burst responses to reward deliveries following probabilistic predictions increased as the probability of reward delivery decreased. During presentations of object sequences, these neurons were most sensitive to the presentation of novel objects and were differentially activated by object sequence violations, though these events were not directly associated with reward. These data show that phasic bursting BF neurons’ activity is most strongly influenced by the surprise and salience of external events.

RPEs which indicate the difference between expected and received rewards, are a crucial variable encoded by many dopamine neurons (Schultz et al., 1997; Cohen et al., 2012), particularly in probabilistic tasks similar to the ones used in this study (Morris et al., 2004; Matsumoto and Hikosaka, 2009; Lak et al., 2014). This dopaminergic value signal may update value or utility functions and directly mediate economic decisions (Schultz, 2010; Lak et al., 2014). However, the data here show that BF phasic bursting activity is unlikely to directly serve the same functions in economic decision. Unlike many dopamine neurons that are suppressed by unexpected reward omissions and are excited by unexpected reward deliveries, BF bursting neurons only signal unexpected reward deliveries. This type of response may be best situated to participate in alerting, learning, and attentional functions, rather than in economic or value-based decisions by signaling when salient and surprising events, such as belief state violations, have occurred.

This hypothesis is further supported by bursting neurons’ responses in the object sequence task, as bursting was greater for novel than familiar objects. Also, a novel object’s task relevance for an upcoming memory-related behavior, increased novelty responses in a gain-like manner. Similarly, object sequence violations that caused sensory surprise, also changed the magnitude of the object-presentation related bursting. These sequence violations did not have an influence on reward rate or expectancy. Most importantly, additional evidence also comes from previous studies that indicate that BF neurons are excited by predictions and deliveries of punishments (Lin and Nicolelis, 2008; Hangya et al., 2015; Monosov et al., 2015).

Indeed, previous work showed that the basal forebrain projects heavily to higher-order prefrontal and temporal sensory areas that process object information and control attention and memory (Mesulam et al., 1983; Everitt and Robbins, 1997; Baxter and Chiba, 1999; Turchi et al., 2005; Lin et al., 2006; Zaborszky et al., 2008; Zaborszky et al., 2013; Lin et al., 2015; Monosov et al., 2015). One recent study showed that inactivation of the BF disrupts the global correlations of these cortical areas with other higher order cortical areas (Turchi et al., 2018). BF can also affect salience and reinforcement timing-related activity even in earlier sensory areas such as V1 (Shuler and Bear, 2006; Lin et al., 2015; Zold and Shuler, 2015). It is therefore likely that on a neuronal circuit level, stimulus-locked BF bursting may coordinate cortical activity in relation to external salient events.

A set of recent studies argue that some dopamine neurons do not signal RPEs wholly or purely (Bromberg-Martin et al., 2010; Matsumoto and Takada, 2013; Takahashi et al., 2016; Takahashi et al., 2017; Babayan et al., 2018). These may complement BF in supporting alerting, attention, and associative learning. Therefore, future studies must assess how the BF and the dopamine system coordinate cognitive functions. An interesting possibility is that BF provides a short-acting/rapid alerting/attentional coordinating signal, correlated with the expectancy or novelty of external events, that is then followed by dopaminergic signaling that may further support associative learning, and economic value-assignments (depending on where the dopamine is released) (Bromberg-Martin et al., 2010; Matsumoto and Takada, 2013; Takahashi et al., 2016; Takahashi et al., 2017; Babayan et al., 2018).

An important consideration for the interpretation of the BF novelty responses is that novelty, in monkeys and humans, is thought to exert strong influence on behavior (Berlyne, 1970; Tiitinen et al., 1994; Bardo et al., 1996; Barto et al., 2013; Foley et al., 2014). Specifically, these studies show that monkeys’ gaze is strongly attracted to novel objects. However, it is unclear whether the motivation driving gaze behavior is mediated by subsequent onsets of novel objects, similar to how this behavior is motivated by subsequent reward deliveries (Hikosaka, 2007). To explicitly test whether novelty can change motivational bias, we designed a new task that directly tested the eagerness of monkeys to saccade to objects that predicted subsequent presentations of familiar or novel objects (Supplemental Figure 10). In that experiment, we found that monkeys were more eager to saccade to familiar objects when, upon target acquisition, the familiar objects were known to be followed by a novel object. Therefore, while BF novelty responses in the object sequence task may not be directly related to primary reward, BF bursting may be related to the motivational effects of novelty on attention and gaze.

A distinct group of BF neurons exhibited tonic and ramping activity and was highly sensitive to the behavioral relevance and timing in our tasks. The tonic activity of these neurons displayed persistent selectivity during the entire 2.5 s CS epoch and was related to the expected values of the CSs. Consistent with previous work (Monosov et al., 2015; Ledbetter et al., 2016), the same neurons displayed ramping activity to the time of uncertain reward delivery or to the delivery of aversive stimuli (Supplemental Figure 9). Here we found that they also ramped to the time of novel stimulus presentation (Supplemental Figure 10) and that their activity is sensitive to both reinforcement deliveries and omissions, signaling the monkeys’ surprise (absolute or unsigned reward prediction errors; Figure 2). This surprise coding was observed after the trial’s outcome, as the neurons ramped down to baseline. Following unexpected omissions in reward-probabilistic trials, in which no sensory cue indicated that reward would not be delivered, the decay of the ramping activity was greatest in trials in which the monkeys were more confident that reward would be omitted (25% CS). The decay rate increased if an explicit sensory cue provided the monkey with a definitive end of the trial (Figure 2). These data show that BF ramping responses signal internal variables closely tied to confidence in the timing of reinforcement delivery.

This observation is also supported in their CS responses. In trials in which there was temporal reward uncertainty, but reward would always be delivered (1.5 or 4.5 seconds after the CS), BF ramping activity was greatest during CS epochs in which reward was most probable. The same neurons’ ramping activity can be elicited by the expectation rewards following long delays (Supplemental Figure 6).

What might be the difference between BF ramping activity and ramping activity observed in the dorsal striatum (White and Monosov, 2016)? First, BF neurons strongly ramp during trace conditioning to the time of the reinforcement. Dorsal striatum uncertainty selective ramping neurons strongly reduced their ramping activations following the disappearance of the CS (White and Monosov, 2016). Second, trial-by-trial variability was very low in the BF ramping neurons (fano factors < 1) while striatal ramping neurons’ trial-by-trial variability was relatively high (Supplemental Figure 11). Finally, ramping activity in the dorsal striatum was specific to reward expectation (White and Monosov, 2016). Therefore, the BF may be particularly well suited to provide an internal clock that is independent of valence and external sensory events.

Because BF is a heterogeneous brain region that contains prominent groups of cholinergic, GABAergic, and glutamatergic projection neurons, the next steps must be to identify which neurotransmitters are released (or co-released) by phasic bursting and ramping neurons. Previous work in rodents has identified putative-GABAergic CS-related phasic bursting neurons, reinforcement salience related bursting cholinergic neurons, and other tonic active neurons in the BF (Lin and Nicolelis, 2008; Unal et al., 2012; Avila and Lin, 2014; Hangya et al., 2015). It will be important to correlate those results from rodents to data recorded in primates. Though species differences in learning of higher-order information (and resulting differences in responses to the violations of these learned states) will likely present significant challenges, such cross-species examinations will be crucial for subsequent unravelling the functional contributions of BF to higher-order behaviors.

## Methods

### General Procedures

Six adult male rhesus monkeys (Monkeys B, R, Z, W, H, and P) were used for recording experiments. All procedures conform to the Guide for the Care and Use of Laboratory Animals and were approved by the Institutional Animal Care and Use Committee at Washington University (Monkeys B, R, and Z) and National Eye Institute (Monkeys P and H). For each monkey, a plastic head holder and recording chamber were fixed to the skull under general anesthesia and sterile conditions. Chambers were tilted laterally from midline by 35 degrees and aimed at the basal forebrain and anterior portion of striatum. After the monkeys recovered from surgery, they participated in behavioral and neurophysiological experiments.

### Data acquisition and analyses

While the monkeys participated in the behavioral procedure we recorded single neurons in the basal forebrain. The recording sites were determined with 1 mm-spacing grid system and with the aid of MR images (3T) obtained along the direction of the recording chamber. This MRI-based estimation of neuron recording locations was aided by custom-built software (Daye et al., 2013). Single-unit recording was performed using glass-coated electrodes (Alpha Omega). During each recording session, an electrode was inserted into the brain through a stainless-steel guide tube and advanced by an oil-driven micromanipulator (MO-97A, Narishige). Signal acquisition (including amplify cation and filtering) was performed using Alpha Omega 44 kHz SNR system. Action potential waveforms were identified online by multiple time-amplitude windows with an additional template matching algorithm (Alpha-Omega).

Neuronal recordings were restricted to single well-isolated neurons in the basal forebrain that displayed task related ramping or phasic-bursting activity following the presentation of the task conditioned stimuli. Basal forebrain was verified as the neuronal tissue within 2 mm relative to the bottom of the brain, since the bottom of the brain can be readily found using electrophysiological criteria. Furthermore, the ventral pallidum (defined using anatomical criteria, such as the first neuronal tissue encountered following traversing the anterior commissure, and previous electrophysiological criteria, such as high and irregular firing rate) was not part of this study because its functions and anatomical projections are distinct from the medial and ventral lateral basal forebrain and because it mostly does not contain the tonic regular firing ramping neurons and phasic bursting low firing neurons.

Eye position was obtained with an infrared video camera (Eyelink, SR Research). Behavioral events and visual stimuli were controlled by Matlab (Mathworks, Natick, MA) with Psychophysics Toolbox extensions. Juice, used as reward, was delivered with a solenoid delivery reward system (CRIST Instruments). Juice-related licking was measured and quantified using previously described methods. Airpuffs were delivered through a narrow tube placed ∼6-8cm from the monkey’s face. Anticipatory licking was acquired and measured using procedures detailed in earlier publications (Monosov and Hikosaka, 2012, 2013).

### Behavioral procedures

#### Probability Amount procedure

To study neuronal representations of reward probability and amount, and delivery related responses following probabilistic reward deliveries, we trained monkeys on a behavioral procedure that contained two blocks of trials: a reward-probability block and a reward-amount block. The trial structure was detailed previously (Monosov and Hikosaka, 2013; Monosov et al., 2015; White and Monosov, 2016). Each trial started with the presentation of a green trial-start cue at the center. The monkeys had to maintain fixation on this trial-start cue for 1 s; then the trial start cue disappeared and one of the CSs was presented pseudo randomly. After 2.5 s (for monkeys B, Z, and R) or 1.5 s (monkeys H and P), the CS disappeared, and juice (if scheduled for that trial) was delivered. The reward-probability block contained five visual fractal object CSs associated with five probabilistic reward predictions (0, 25, 50, 75 and 100% of 0.25 ml of juice). The reward-amount block contained five objects associated with certain reward predictions of varying reward amounts (0.25, 0.1875, 0.125, 0.065 and 0ml). Each block consisted of 20 trials (for monkeys B, Z, and R) and 40 trials (for monkeys P and H) with fixed proportions of trial types (each of the five CSs appears four times in each block).

Before neuronal recordings began, the monkeys’ knowledge of the CSs was confirmed by a choice procedure, and the choice data was previously published (Monosov et al., 2015; Monosov, 2017). Briefly, in separate experimental sessions, the monkeys’ choice preference was tested for the CSs. Each trial started with the presentation of the trial-start cue at the center, and the monkeys had to fixate it. Then two CSs appeared 10 degrees to the left and right. The monkeys had to make a saccade to one of the two CSs within 5 s and fixate it for at least 750 ms. Then the unchosen CS disappeared, and after a brief delay the outcome (associated with the chosen CS) was delivered, and the chosen CS disappeared. If the monkey failed to fixate one of the CSs, the trial was aborted and all stimuli disappeared. The trials were presented pseudo randomly, so that a block of 180 trials contained all possible combinations of the 10 CSs four times. To verify that the monkeys’ knowledge is stable during recording, we also monitor licking behavior and confirm that it, like the choices, is correlated to the expected values of the probability CSs and amount CSs (two separate Spearman’s correlations, threshold: p<0.05). The CS epoch responses of the 31 neurons recorded in Monkeys H and P were previously analyzed in Monosov et al., 2015.

#### Temporal uncertainty procedure

To assess how monkeys’ BF neurons encoded uncertain predictions about reward timing, monkeys B, R, Z were trained on an additional Pavlovian procedure (Supplemental Figure 4). Following a trial start cue fixation period (same as above), one of five CSs were presented. These CSs predicted either (1) a probabilistic delay before a reward with deterministic delivery (reward-timing-uncertain CSs); or (2) a deterministic delay before a reward with some probability of delivery (reward-probability CS). In trials with one of the four reward-timing-uncertain CSs, reward was always delivered either 1.5 s after CS onset or 4.5 s after CS onset. Depending on the reward-timing uncertain CS, the probability that a reward was delivered at 1.5 s with 0.25, 0.50, 0.75, or 1 probability. In trials with the reward-probability CS, reward was delivered with a delay of 1.5 s after CS onset with 0.50 probability. During, the 0.25, 0.50, and 0.75 CS trials, when reward was not delivered at 1.5 s, the CS remained on the screen until reward was delivered at 4.5 s. During the 0.50 reward probability CS, the CS turned off at the time of the outcome (when reward was either delivered or omitted).

#### Object sequence procedure

An object sequence task was used to study how BF neurons encode sensory predictions and object novelty. Monkeys B, R, and Z experienced four distinct sequences of object presentations (S1, S2, S3, S4). The object sequences began following a 0.5 s period of fixation on the trial start cue that appeared in the center of the screen. Each sequence contained 3 familiar objects and 1 novel object. These objects were presented in the center of the screen and occupied a |3 degree visual angle. The novel object was always presented in second position in the sequence. Therefore, the novel object was surprising because it was never experienced by the monkeys, but its presentation did not deviate from the animals’ expectations. Monkeys performed more than 10,000 trials before recordings began. Following sequences S2 and S4, the monkeys performed a reaction-time Delayed Non-matching-to-Sample task (DNMS). During DNMS, an object that was novel during the presentation of S2 (or S4 if the DNMS trial followed S4) was presented with a novel object that has never been experienced. The objects were presented 10 degrees from the center, to the left and the right of the fixation point. The trial continued until the monkeys fixated the novel object for 0.5 milliseconds to get a reward. The monkeys were never penalized for looking at the previously experienced object. Therefore, the novel objects in S2 or S4 did not have an explicit reward association, but aided the monkey in subsequent DNMS trials. On |11% of S2 or S4 presentations, the first or the third fractal was replaced by a corresponding fractal from sequences S1 and S3 (in S2 from S1; and in S4 from S3). For example, if the first fractal in S2 was replaced, the first fractal from S1 was always displayed. In this way, sequence violations did not alter the relationship of the individual fractals to the timing of reward delivery.

#### Reward and novelty motivated gaze task

To test if monkeys are motivated by novelty we trained Monkeys R and Z on a saccadic task that measured their eagerness to observe a novel visual object. First, a fixation dot appeared in the center of the screen. 0.5 s after the onset of the fixation dot, a visual object fractal appeared 10 degrees to the right or the left of the fixation dot. The monkey was required to continue fixating the dot in the center. After 0.35 s the fixation spot disappeared and the monkey was free to make saccades. Reward was always delivered 3 seconds after the fractal onset. Therefore, the monkeys’ saccadic behavior after the fixation spot disappeared did not affect reward delivery. In this task, the monkeys experienced four different trial types. The first two types of trials contained a novel (type 1) or 1 of 2 familiar (type 2) visual fractal objects. Two additional trial types (3-4) tested whether the monkeys were motivated by the possibility of viewing a novel fractal. In trial type 3, 1 of 2 familiar objects appeared. After fixation spot disappeared, if the monkey fixated the familiar object, it was immediately replaced by a novel object. In trial type 4, 1 of 2 familiar objects appeared. If the monkey fixated this object, it was replaced by 1 of 2 familiar objects. If novelty is salient, we ought to observe faster target acquisition times (duration between the time when the stimulus was presented and when the monkey saccades to its location) in trial type 1 than 2. Also, if novelty exerts motivational effects on saccadic behavior, then we ought to see faster target acquisition times in trial type 3 than 4.

### Analyses

To generate spike density functions, spike times were convolved with a Gaussian kernel (σ=100 ms) across all trials. Statistical tests were two-tailed. All permutation tests used 10000 shuffles. For all analyses and figures that included deliveries and omissions of rewards, unless explicitly stated in the text, a neuron was included if it had at least 2 trials for reward delivery and omission.

To cluster the single neurons’ average responses in the probability block (Figure 1), first we performed principal component analysis (PCA). We then applied Silhouette and Calinski-Harabasz tests to estimate the optimal number of clusters (n=2). K-means clustering was used to cluster the data based on PCs into 2 clusters (for this, using the first 3 PCs and up to 10 PCs resulted in very similar group membership).

To calculate the latency of reward size coding information (Figure 1C) we performed a correlation of firing rate and value in time (in 100ms bins moving 1 ms steps) for each neuron.

For each time bin we calculated the p value of the Spearman’s rank correlation of neuron’s activity with reward amount in the reward amount block. Reward size coding latency was defined as the first time p was lower than 0.01 (but similar results were obtained at p < 0.05). Importantly, such statistical latency analyses do not determine the actual latency of information coding in the brain because they utilize an arbitrary threshold. Instead, they are useful for demonstrating relative latencies across two groups of neurons or signals (Monosov et al., 2008).

To calculate the baseline rate that was used to derive the latency with which ramping neurons returned to baseline (Figure 2E), we picked the time window from 1000ms to 500ms before trial start cue appeared and used the average firing rate in this time window as the baseline.

To fit the outcome related activity with exponential functions (Figure 2), we first derived spike density functions using overlapping bins of 100ms (in 50ms steps). We fit the function: *A*\*e^λt^+*C*, in which is the decay rate, representing how fast the firing rate decreases. To determine if the decay rates were significantly different across the different reward-omission conditions we utilized bootstrapping to calculate the confidence interval of the difference between the two decay rates and tested if the 99% or 95% confidence interval excluded a difference of zero. Bootstrapping was done by randomly resampling the neurons with replacement (500 times); each time resampling was done, we obtained a set of decay rates by fitting the neurons’ average activity to the function shown above. In Figures 2A-D, data from probability-amount and reward timing procedures were pooled because the outcome related activity did not differ across them (see Supplemental Figures 1 and 5).

In the DNMS object sequence task, reward was delivered as long as the monkey fixated on the novel object for 0.5 s, regardless if he had looked at the other object. To evaluate the monkey’s performance, we focused on the primary choice the monkey made, i.e., the first object he fixated for 0.5 s. To calculate performance, we obtained the percentage of trials in which the monkeys’ primary choices were the novel objects.

For single neuron analyses (Figure 4B-E) of novelty, task-relevance, and sequence-violations in the object sequence task, we subtracted the activity 100 ms before the object presentation from the activity measured after the object was presented (time window: 200ms to 400ms). In this way, changes in firing rate that were unrelated to the objects were not considered in the analyses.

Neuronal discrimination of object novelty was assessed by calculating area under the receiver operating characteristic (ROC) curve. ROC areas of 0 and 1 are equivalent statistically; both indicate that two distributions are completely separated. The analysis was structured so that ROC area values greater than 0.5 indicate that the activity during novel object presentation was greater than familiar.

To measure spike time variability across trials we derived the Fano factor (FF). FF was calculated in 100ms overlapping bins (in 1 ms steps).

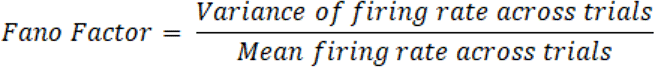

## Supplemental Figure Legends

**Supplemental Figure 1.**
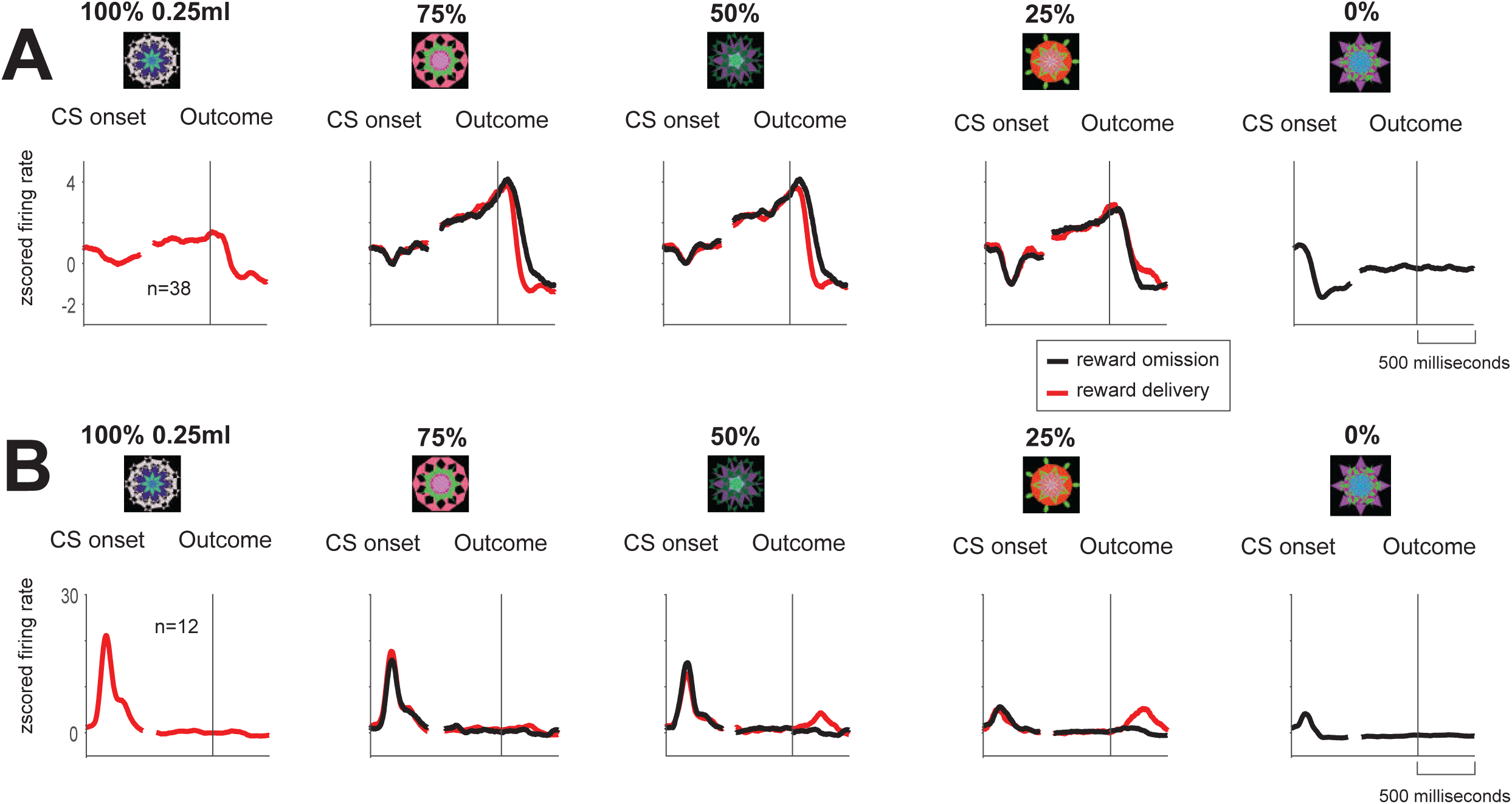
BF activity across five probabilistic reward predictions. Neurons’ average activity shown separately in trials in which rewards were predicted with 5 different reward probabilities (indicated on the top; actual fractals used in the task are shown above the neuronal activity). After the trials’ outcome time, activity is shown separately for reward delivered trials (red) and reward omitted trials (black). (**A**) Ramping neurons (**B**) Phasic bursting neurons. In this figure neurons with at least 2 trials for each condition (e.g. delivery versus omission) are shown.

**Supplemental Figure 2:**
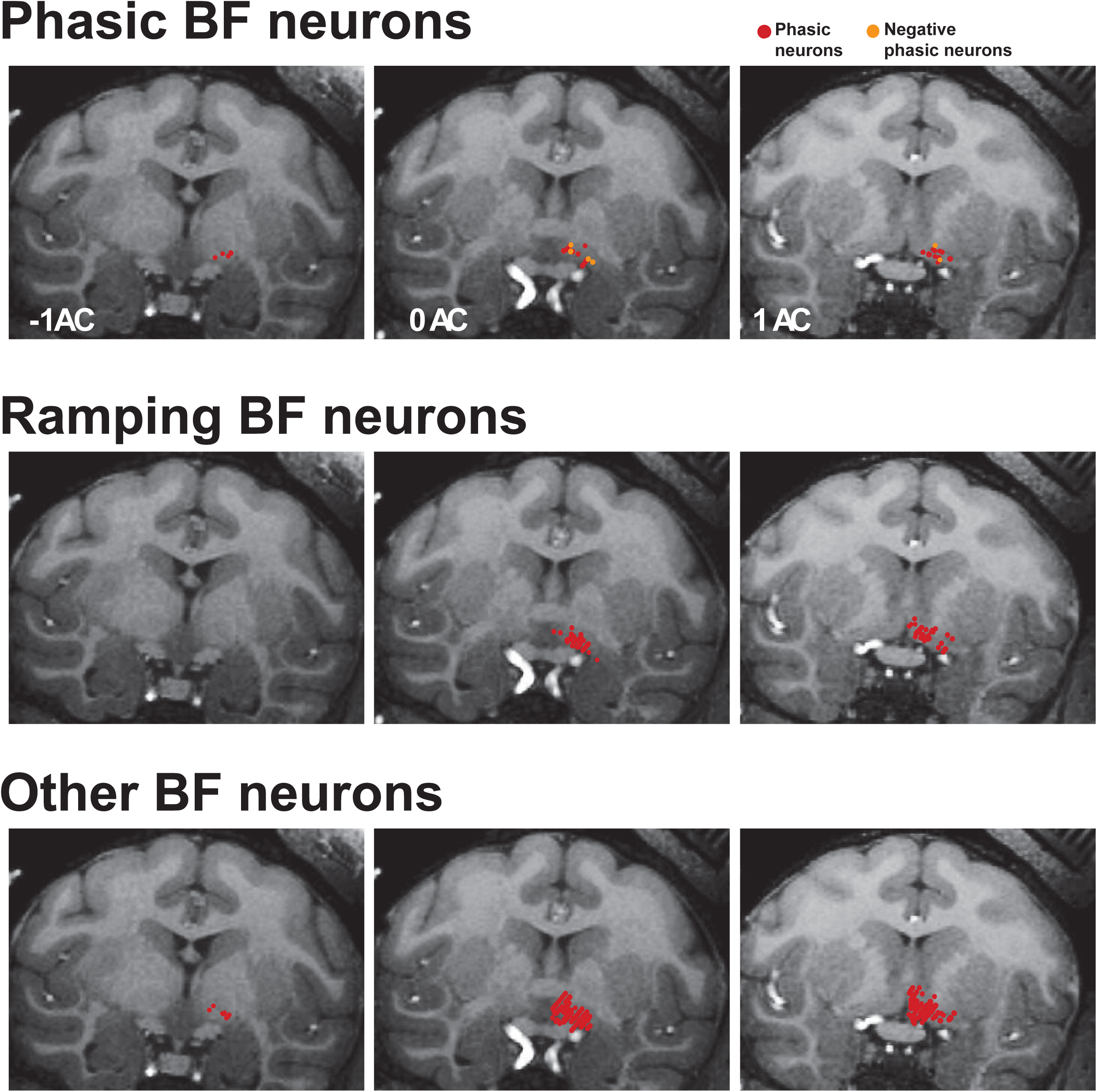
Estimated locations of phasic bursting and tonic ramping neurons in the BF. The recording range in the BF was -2 to 4 mm anterior to the center of the anterior commissure (AC). Phasic neurons (top; n=38), tonic ramping neurons (middle; n=79), and other neurons encountered online that did not have ramping or phasic activity (n=280; bottom) are shown on three coronal T1 MRI images. All single neurons across all tasks are shown here. Because some neurons were recorded in 1 or several tasks the neuron number here is greater than in Figure 1.

**Supplemental Figure 3:**
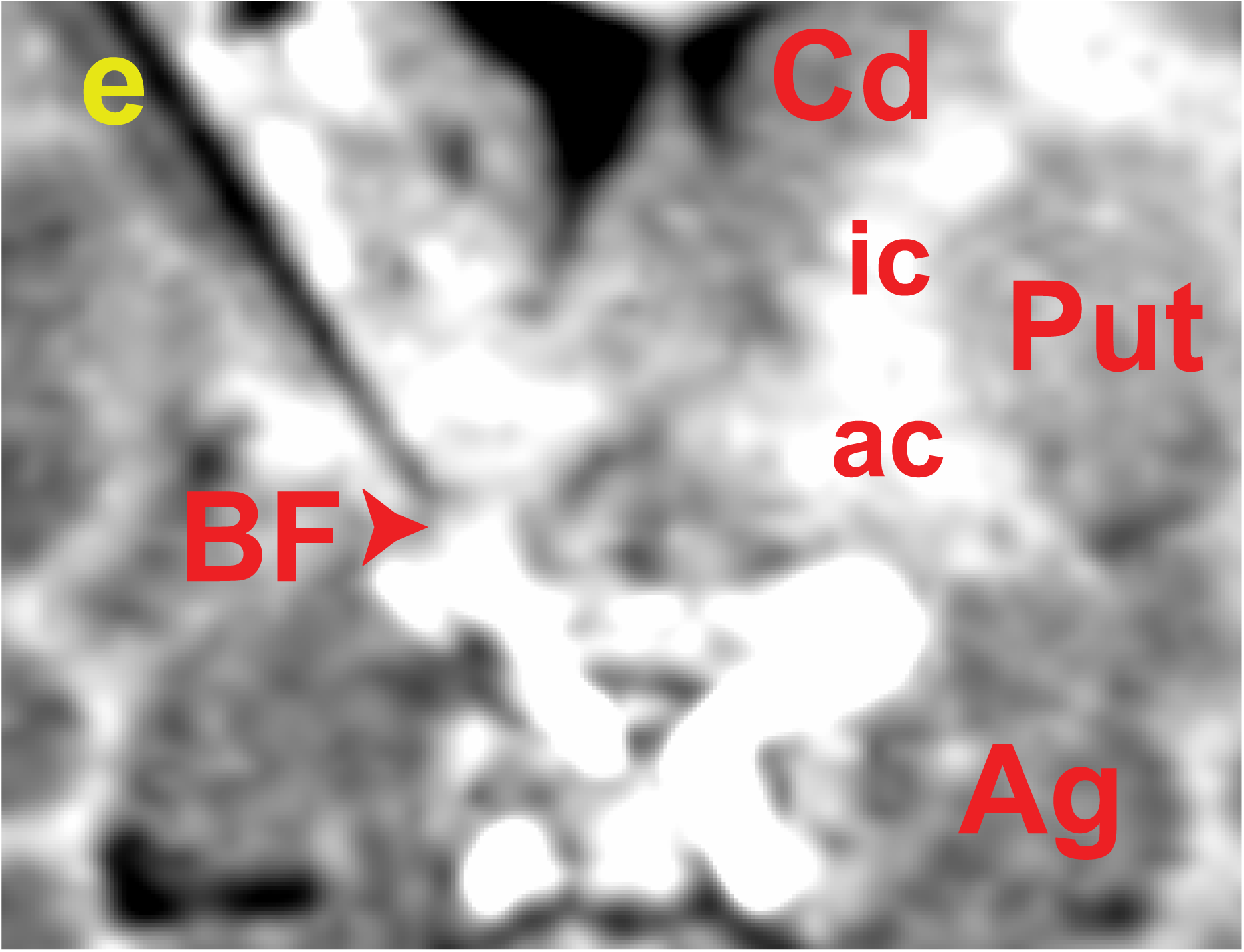
Confirmation of recording location within BF. A coronal MRI confirming a recording location of a phasic bursting neuron within the BF of monkey B. The image was acquired with a tungsten electrode (FHC) at the recording location within BF. The electrode’s shadow is the black line whose tip is in BF (marked by a yellow **e**). BF – basal forebrain; ac – anterior commissure; ic – internal capsule; Cd – caudate; Put – putamen, Ag – amygdala.

**Supplemental Figure 4:**
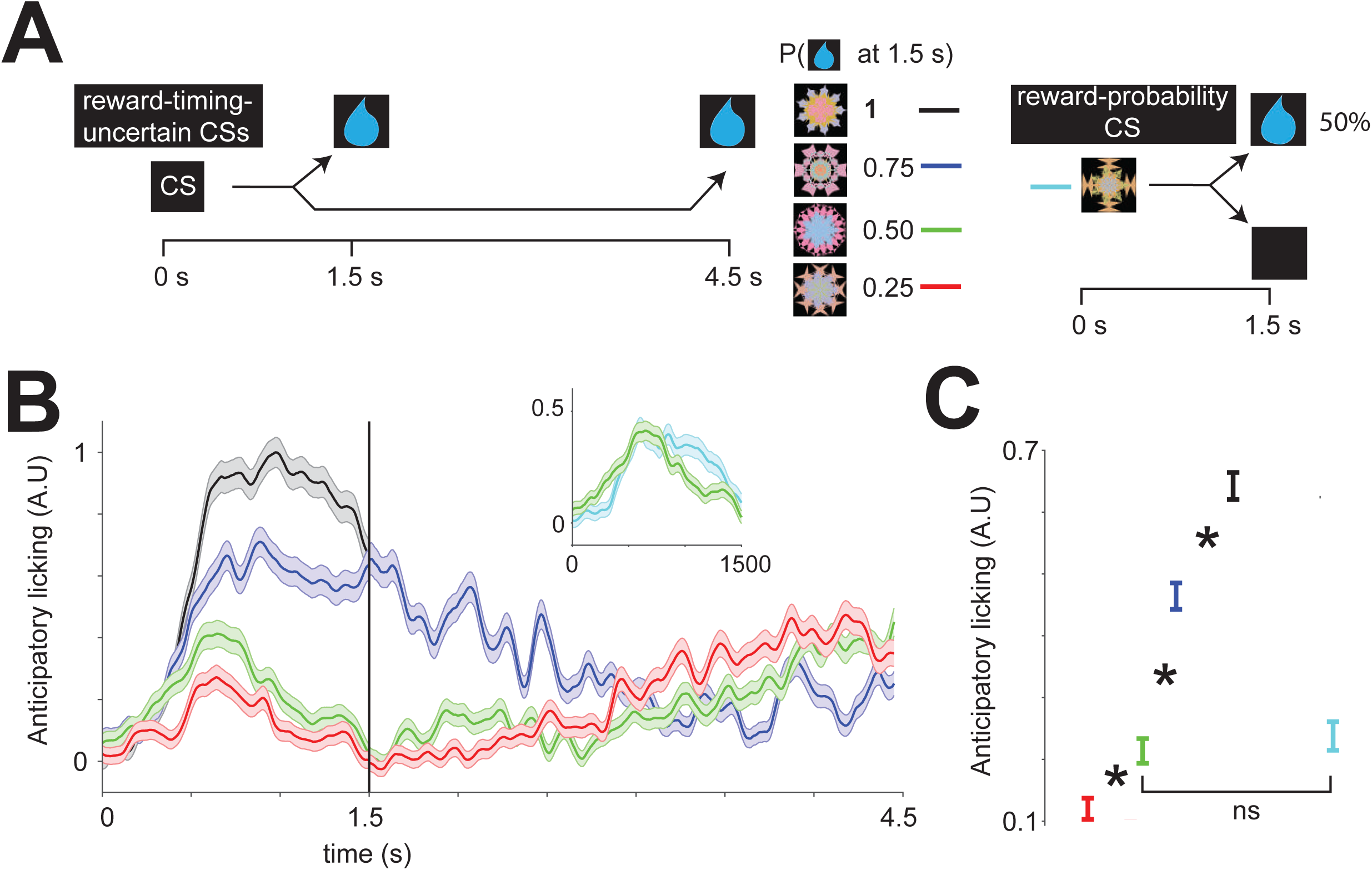
Reward timing task. **(A)** The reward timing procedure uses Pavlovian delay conditioning in which five distinct visual fractal objects serve as conditioned stimuli (CSs) that predict either (1) a probabilistic delay before a reward with deterministic delivery (reward-timing-uncertain CSs); or (2) a deterministic delay before a reward with some probability of delivery (reward-probability CS). In trials with one of the four reward-timing-uncertain CSs, reward is delivered at the latest with a delay of 4.5 s after CS onset. However, depending on the CS, reward has either a 0.25 (red), 0.50 (green), 0.75 (blue), or 1 (black) probability of being delivered earlier with a delay of 1.5 s after CS onset. In reward-probability trials, reward is delivered with a delay of 1.5 s after CS onset with 0.50 (cyan) probability. **(B)** Time course of anticipatory licking behavior is shown before possible reward delivery at 1.5 s across all trials; and from 1.5 s to 4.5 s across trials with a reward-timing-uncertain CS in which reward was not delivered at 1.5 s. **(C)** The mean magnitude of anticipatory licking behavior increases with the probability of reward delivery at 1.5 s (Spearman’s rank correlation, ρ=0.38, p<0.0001). The asterisks indicate significant differences between CSs (Wilcoxon rank-sum test, p<0.05). The ‘ns’ indicates no significant difference between CSs (Wilcoxon rank-sum test, p>0.05).

**Supplemental Figure 5:**
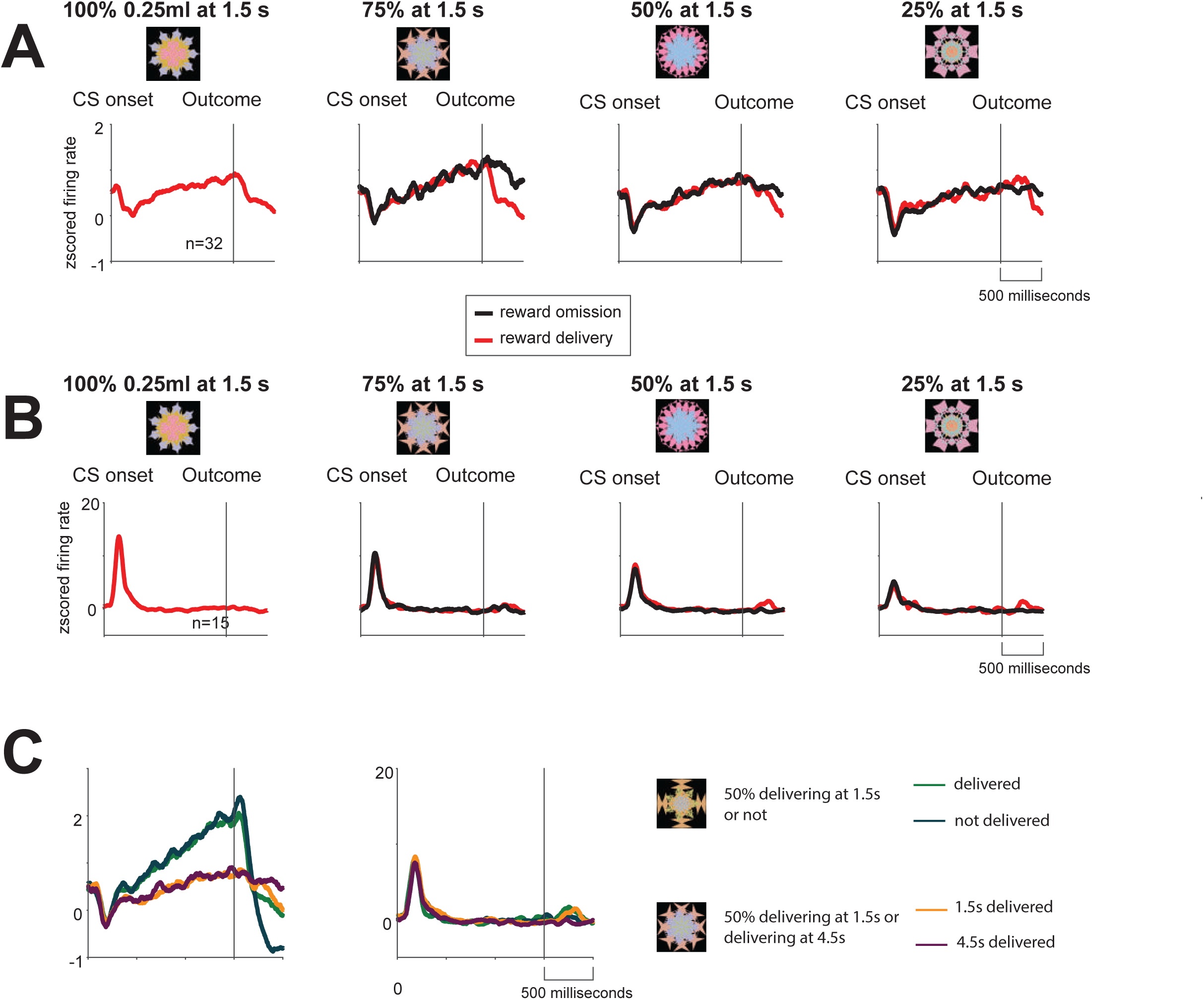
BF activity across reward timing-uncertain trials. **(A)** Tonic ramping neurons’ average activity shown separately in trials in which rewards were predicted at 1.5 s with 4 different probabilities (indicated on the top; actual fractals used in the task are shown above the neuronal activity). After the trials’ outcome time, activity is shown separately for reward delivered trials (red) and reward omitted trials (black). (**B**) Phasic bursting neurons. (**C**) The CS related activity of tonic ramping (right) and phasic (left) BF neurons during 50% reward trials in which reward was delivered or omitted at 1.5 s and 50% reward trials in which reward was delivered at 1.5 s (50% of the trials) or 4.5 s (if it was omitted at 1.5 s). Like the licking behavior (Supplemental Figure 4) phasic bursting neurons did not discriminate between these two types of trials (p = 0.51, sign rank test, time window: 100ms to 600ms after fractal onset, the data of delivering or not at 1.5s had been combined, same as delivering at 1.5s or 4.5s. The combination was also applied to ramping neurons). Tonic ramping neurons displayed stronger ramping to the 50% reward probability CS that was riskier (p <0.01, time window: 1000ms to 1500ms after fractal onset, sign rank test). In A-C, neurons with at least 2 trials for each condition (e.g. delivery versus omission) are shown.

**Supplemental Figure 6:**
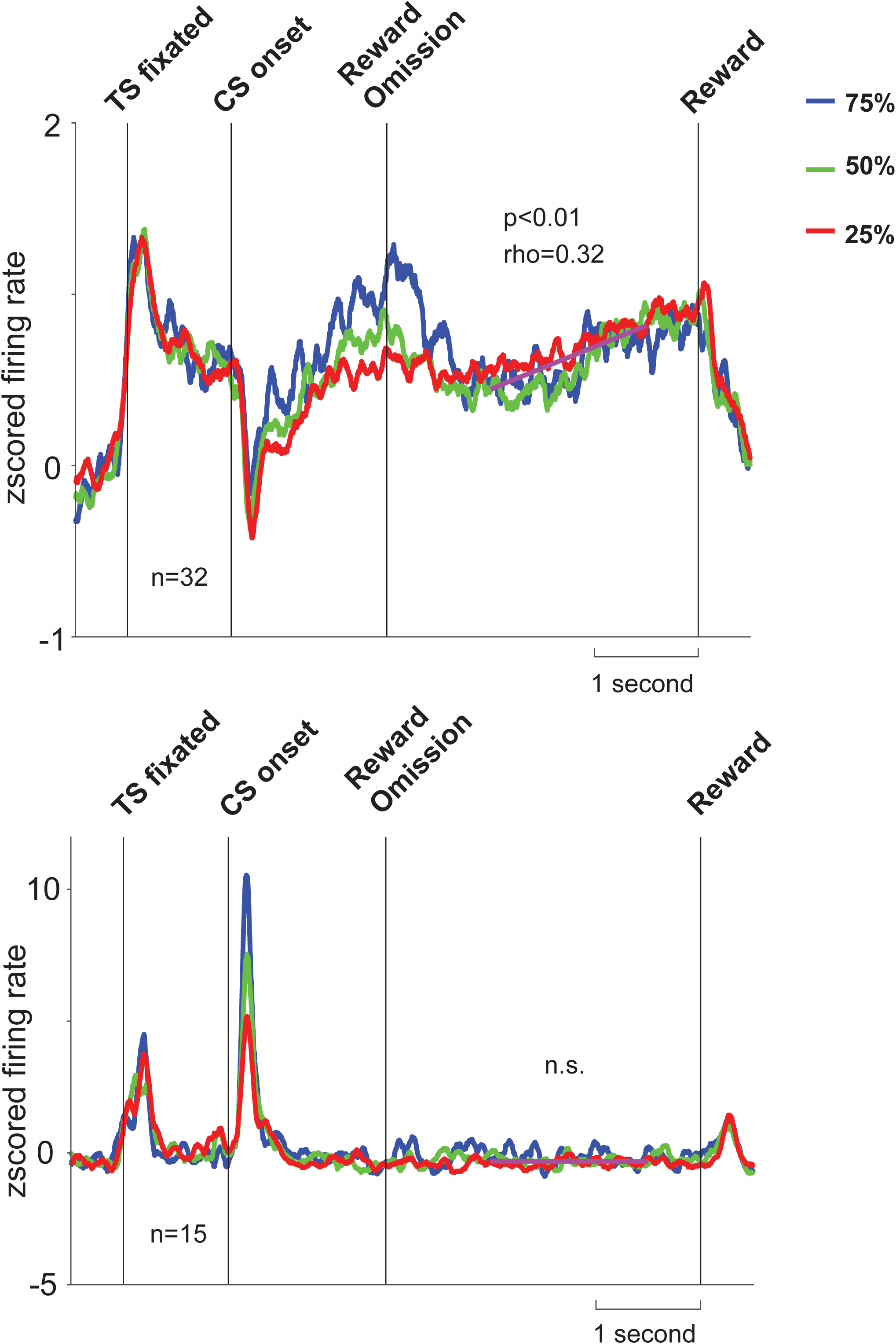
BF activity across reward timing-uncertain trials after reward omissions. Reward-timing-uncertain CS responses and post-outcome reward-timing signals. Mean neuronal activity of the tonic ramping neurons **(top)** and phasic bursting neurons **(bottom)**. Activity from all trials is shown before 1.5 s, and only from trials in which rewards were not delivered at 1.5 s is shown after 1.5 s. After reward was not delivered at 1.5 s, tonic ramping neurons displayed anticipatory ramping activity for the 4.5 s reward (average ramping across all 3 conditions: Spearman’s rank correlation, ρ=0.32, p<0.01; time window: 2 seconds before 4.5s reward delivery), while phasic neurons’ activity have a much weaker ramping, p=0.6.

**Supplemental Figure 7:**
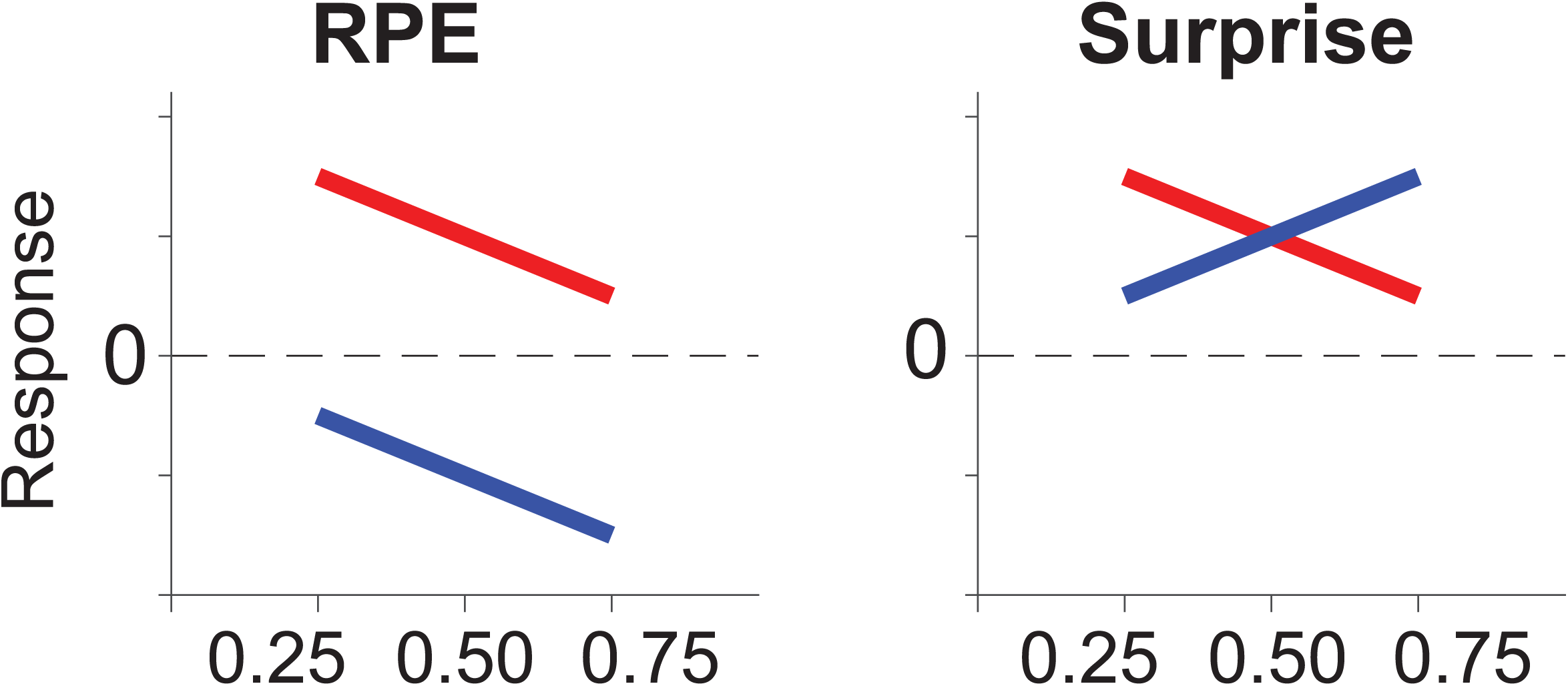
Toy models of RPE and surprise. Cartoon models of reward prediction error coding (left) and reward surprise coding (right). Theoretical responses to reward deliveries (red) and omissions (blue) following different reward probabilities (x-axis).

**Supplemental Figure 8:**
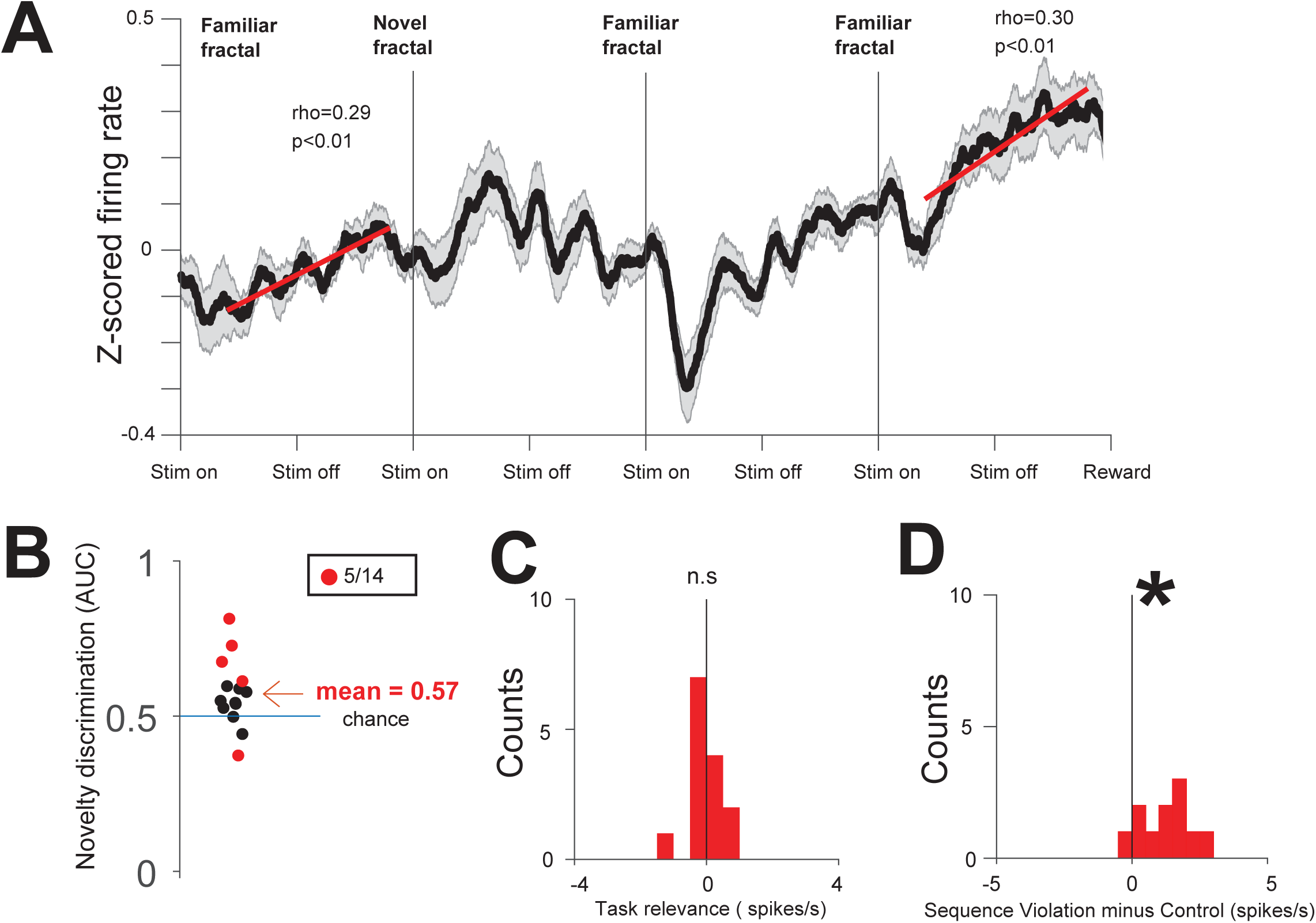
Tonic ramping neurons’ responses in the object sequence task. (**A**) Average activity of ramping neurons in the object sequence task. (**B**) Area under the ROC curve (AUC) for each ramping neuron that assessed the ability of the neuron to discriminate novel versus familiar objects. Red dots neurons that significantly discriminate novel versus familiar objects (time window: 200 ms to 400 ms). (**C**) Histogram of single neurons’ response differences for novel fractal in Sequence 2 and Sequence 1, there is no significant difference from 0 (p = 0.79). (**D**) Same convention as Figure 4E. Here the object sequence violation also significantly enhanced neurons’ responses (p<0.01).

**Supplemental Figure 9:**
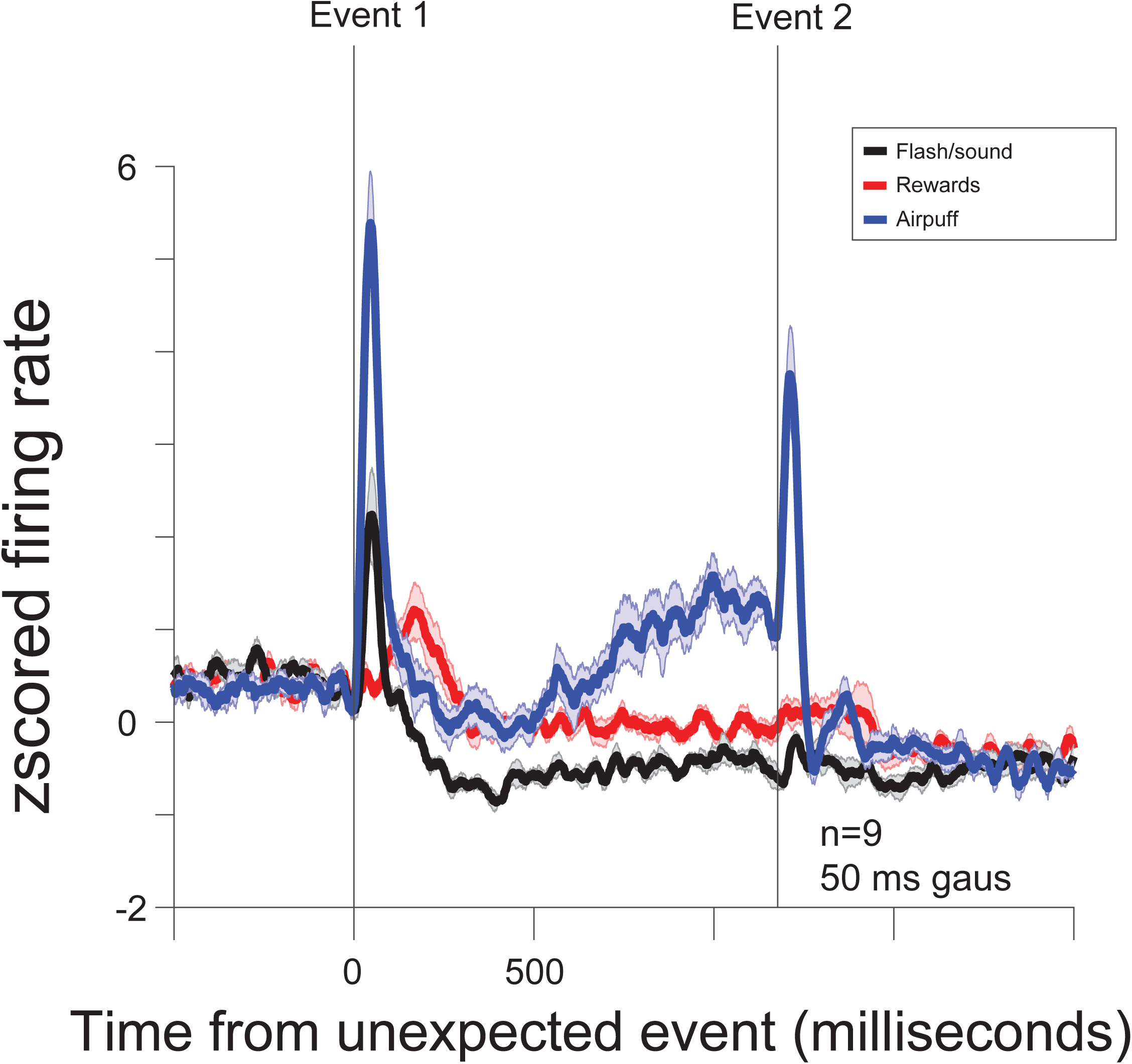
BF tonic ramping neurons’ responses to salient stimuli. In the object sequence task, we found that tonic ramping neurons display ramping activity to the time of the novel stimulus. Might ramping occur in anticipation of any salient stimulus or in anticipation of a distracting stimulus? This question is important because the same class of neurons has been shown to ramp to the time of punishment deliveries, and respond phasically to the delivery of unexpected rewards and punishments during the inter-trial-interval. To answer the question, we trained Monkey W to participate in a simple Pavlovian procedure with two CSs associated with two probabilities of reward (100 and 50%). The key was that during |15% of the inter-trial-intervals, between the outcome and the next trial’s start cue, we presented an unexpected 150 ms event comprising of a color change on the screen (while luminance was fixed) and a synchronous white noise burst. If this event occurred, a second audiovisual event occurred after a fixed time interval. As expected BF ramping neurons responded phasically to the first event (black trace). However, after this, they did not ramp to the second event. In fact, their activity was clearly reduced relative to baseline. To replicate previous work, we also included double rewards (red) and double air puff punishments (blue) (such that any of the 3 types of events occurred with equal probability). As expected, during double rewards, the neurons responded to the first event with a phasic burst, and returned to baseline; and as shown previously, both unexpected and expected punishments elicited a response. Also, the neurons ramped to the time of punishments (as previously shown in Monosov, et al., 2015). To summarize, is that the BF ramping neurons do not ramp to the time of irrelevant events (here an audio-visual event). In fact, they reduced their baseline activity when irrelevant events were expected. To clearly visualize the rapid phasic events, here we used 50 ms Gaussian kernel for to generate the spike density functions. Shaded regions represent SEM.

**Supplemental Figure 10:**
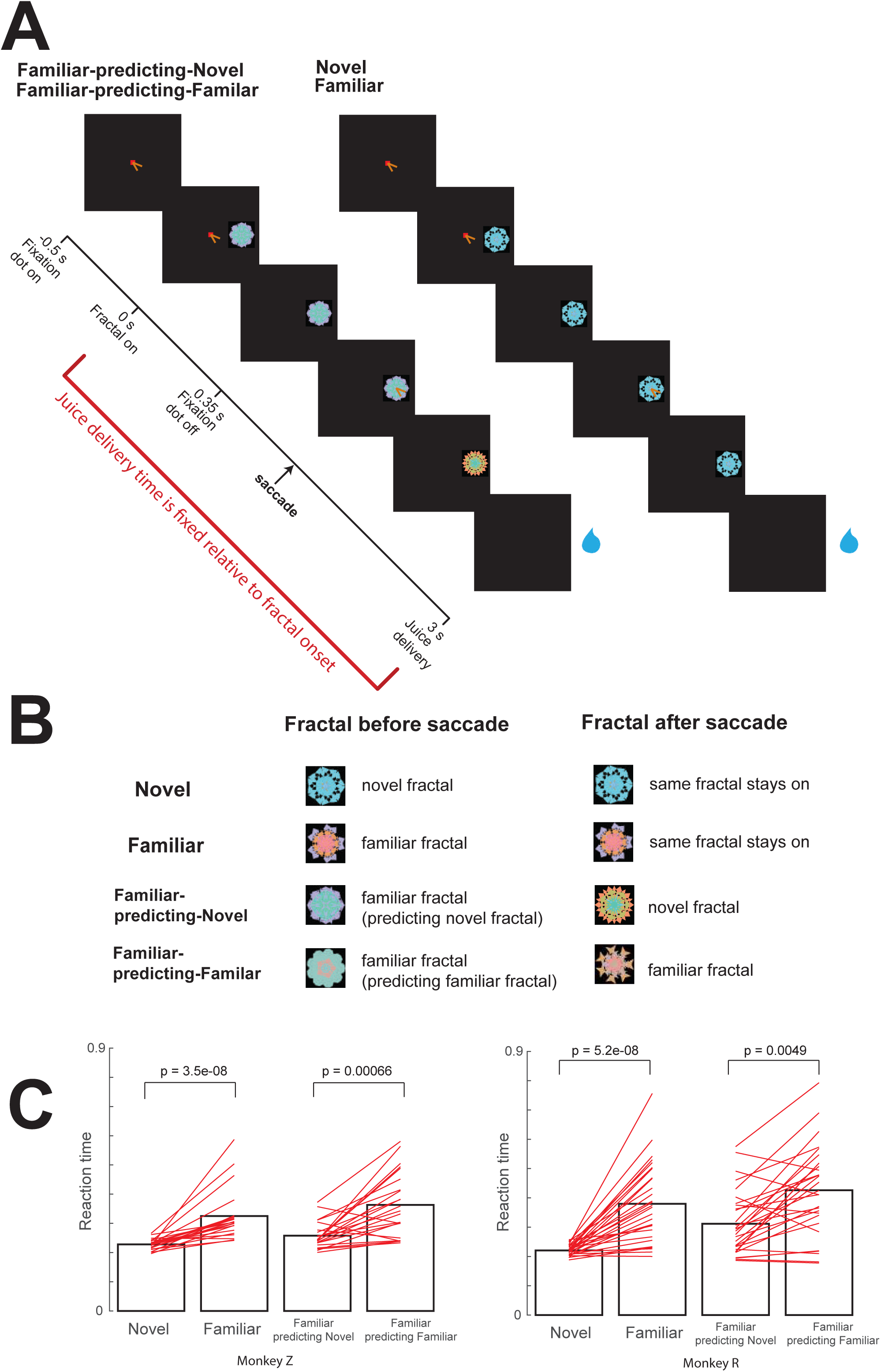
Motivational- and sensory-salience effects of novelty on gaze behavior. (**A**) To test if monkeys really are behaviorally motivated by novelty we trained Monkeys R and Z on a novel saccadic task that measured their eagerness to observe a novel visual object. First, a fixation dot appeared in the center of the screen. 0.5 s after the onset of the fixation dot, a visual object fractal appeared either to the right or the left of the fixation dot (angle: 10 degrees). The monkey was required to continue fixating the dot in the center. After 0.35 s the fixation spot disappeared and the monkey was free to make saccades. Reward was always delivered 3 seconds after the fractal onset. Therefore, the monkeys’ saccadic behavior after the fixation spot disappeared did not affect reward delivery. (**B**) Monkeys experienced four different trial types. The first two types of trials contained a novel (type 1) or 1 of 2 familiar (type 2) visual fractal objects. Two additional trial types (3-4) tested whether the monkeys were motivated by the possibility of viewing a novel fractal. In trial type 3, 1 of 2 familiar objects appeared. After fixation spot disappeared, if the monkey fixated the familiar object, it was immediately replaced by a novel object. In trial type 4, 1 of 2 familiar objects appeared. If subsequently the monkey fixated this object, it was replaced by 1 of 2 familiar objects. (**C**) After training (8 days for Monkey Z and 5 days for Monkey R), Monkey Z (left) and Monkey R (right) displayed a decrease in target acquisition reaction time (the time from fixation off to the time the eye fixates the peripheral object) during trials in which the peripheral object was novel versus familiar (first versus second bar), and when a familiar peripheral object was associated with the presentation of a novel object (third versus fourth bar). Bars indicate mean target acquisition times across all sessions, and the single lines are single sessions’ means.

**Supplemental Figure 11:**
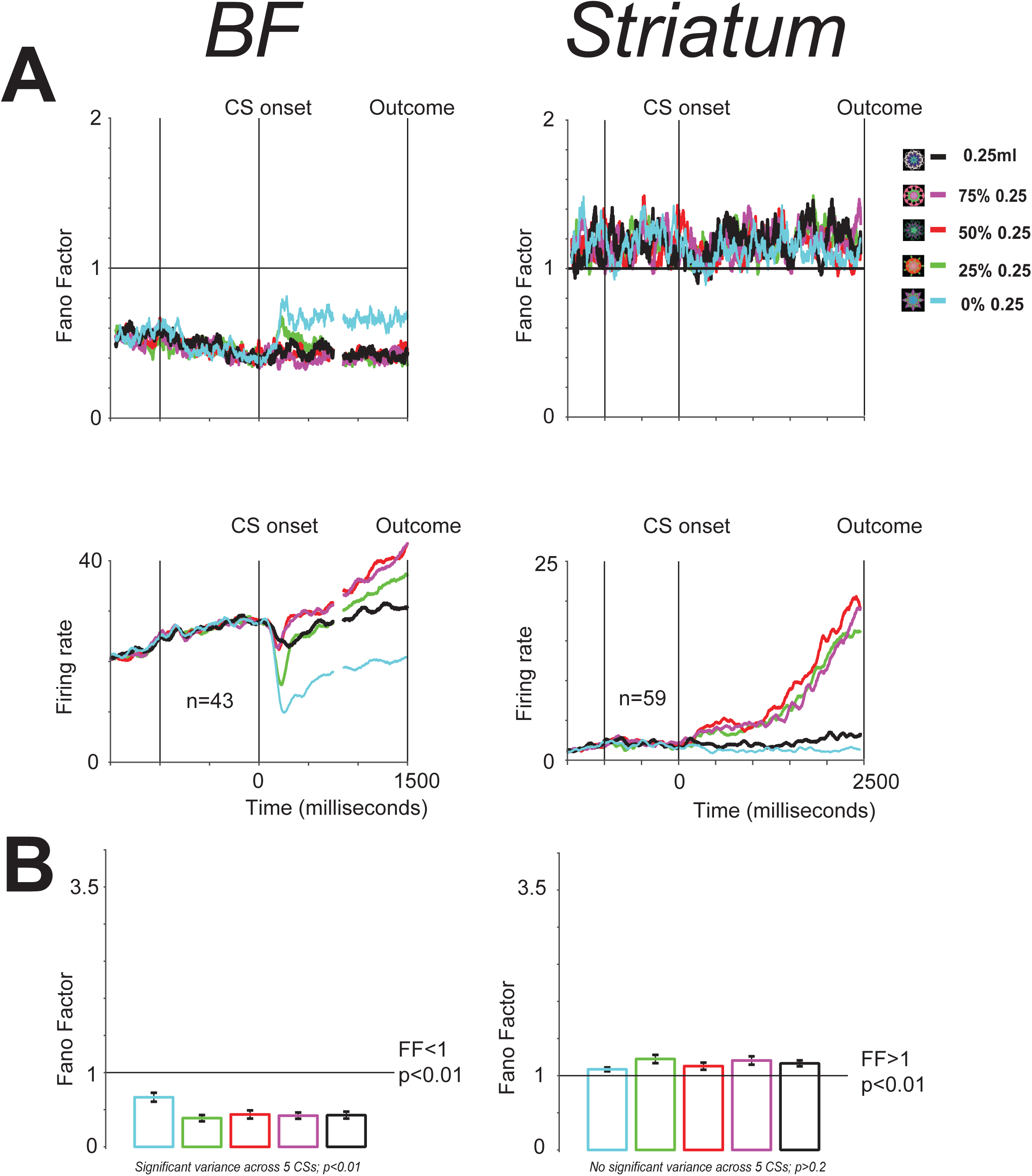
BF ramping neuron’s activity has low trial to trial variability. (**A**) Fano factors (FF) for BF ramping neurons (left) and for comparison striatal uncertainty ramping neurons (right). Their CS related average population activity is shown below (left – BF ramping neurons; right – striatal ramping neurons). (**B**) Population average FF for each 5 CS conditions in the reward the reward probability block for BF ramping neurons (left) and striatal ramping neurons (right). In the BF, FF is lower than 1, indicating low trial to trial variability (less than Poisson). In the striatum, FFs were higher than 1 (and higher than in BF; two rank sum tests; each resulting in p<0.01). Finally, in the BF FF was not directly related to firing rates. In general, BF FF was highest during no reward CS conditions (p<0.01, rank-sum test, comparing no reward conditions with all other conditions), and similar across all other CSs (though firing rates were markedly different, p = 0.8, Kruskal-Wallis test). In contrast, striatum had similar FF across all 5 CS conditions.

## Acknowledgements

This work was supported by the National Institute of Mental Health under Award Number R01MH110594 and the Defense Advanced Research Projects Agency (DARPA) Biological Technologies Office (BTO) ElectRx program under the auspices of Drs. Eric Van Gieson and Doug Weber through the CMO Grant/Contract No.HR0011-16-2-0022. We are grateful to Ms. Julia Pai, Dr. Noah Ledbetter and Mr. J. Kael White for assisting in data acquisition, to Ms. Kim Kocher for fantastic animal care and training, and to Dr. Okihide Hikosaka for allowing us to use data from Monkeys H and P. We thank Ms. Julia Pai and Dr. Ethan Bromberg-Martin for detailed discussion of analyses methods, and Dr. Hiroyuki Nakahara and Ms. Jamie Moffa for reading earlier versions of this manuscript.

